# Training background shapes the exercise-induced extracellular vesicle proteome to acute exercise with partial overlap with cold exposure in humans

**DOI:** 10.64898/2026.11.19.767541

**Authors:** Dominik Tischer, Alexander Braunsperger, Sebastian Kallabis, Daniela Schranner, Paul Jonas Jost, Jan Hasenauer, Felix Meissner, Alexander Pfeifer, Henning Wackerhage, Ana Soriano-Arroquia

**Affiliations:** Institute of Pharmacology and Toxicology, Biomedical Center, University Hospital, University of Bonn, Venusberg Campus 1, 53105 Bonn, Germany; Exercise Biology Group, Department of Sport and Health Science, TUM School of Medicine and Health, Technical University of Munich, Am Olympiacampus 12, 80809, Munich, Germany; Institute of Innate Immunity, Department for Proteomics and Signal Transduction, Medical Faculty, University of Bonn, Bonn, Germany; Core Facility Translational Proteomics, Institute of Innate Immunity, Medical Faculty, University of Bonn, Bonn Germany; Institute of Computational Biology, Helmholtz Zentrum München, Ingolstädter Landstraße 1, 85764 Neuherberg, Germany; Computational Life Sciences, Life and Medical Sciences (LIMES) Institute and Bonn Center for Mathematical Life Sciences, University of Bonn, Bonn, Germany

**Keywords:** extracellular vesicles, exercise, cold exposure, proteomics, training status

## Abstract

**Aim:** Exercise and cold exposure changes circulating extracellular vesicle (EV) and modifies their molecular cargo. However, it remains unclear whether the acute exercise-induced EV proteome response is uniform or shaped by long-term training background, and to what extent it overlaps with the response to cold exposure. We therefore performed an exploratory proteomics study in humans with distinct training phenotypes and compared exercise-induced protein changes with an independent cold exposure dataset.

**Methods:** Serum samples were collected before and immediately after a maximal bicycle ramp test from untrained sedentary controls, endurance-trained athletes, natural bodybuilders, and sprinters. For cold exposure studies, serum samples were collected before, immediately after, and 30 minutes after an ice bath (8-12 °C) for 5 minutes. EVs were isolated by differential ultracentrifugation and characterized using electron microscopy and nanoparticle tracking analysis. EV proteomics was analyzed by LC-MS/MS. Differential relative protein abundance was assessed within each training group. Shared significance, concordant directionality, functional enrichment, and comparison with the independent cold-water immersion cohort were analysed.

**Results:** Acute exercise altered the serum EV-enriched proteome in all training groups, but the response differed substantially between groups. Sprinters showed the largest number of altered proteins, whereas endurance athletes, bodybuilders, and sedentary controls displayed more selective responses. Few proteins reached nominal significance across multiple groups, although a broader set showed concordant directional changes, indicating a limited shared response superimposed on training background-specific remodeling. Functional enrichment highlighted extracellular vesicle and extracellular compartment signatures together with immune, adhesion, trafficking, and stress-response pathways. Cold exposure also induced time-dependent remodeling of the serum EV-enriched proteome, with the largest number of altered proteins observed 30 minutes after exposure. Comparison of the immediate cold response with exercise revealed partial directional overlap, particularly with endurance exercise, but also clear stimulus-specific differences.

**Conclusion:** Our results suggest that changes in the EV proteome profiling after acute exercise are highly dependent on the training background in humans. Rather than revealing a single universal EV protein signature, acute exercise produced group-specific EV proteome remodeling together with a smaller layer of shared directional trends. The comparison with cold exposure supports the concept that acute physiological stressors can remodel concomitant circulating EV protein profiles, while also producing stimulus-specific signatures.

**Highlights:**

- Acute exercise remodeled the EV proteome in a training-background-dependent manner.
- Sprinters showed the largest acute EV proteome response.
- No overlap of significant proteins between different training status.
- Proteome profiles show concomitant directionally trends among all training groups.
- Cold exposure shows specific EV-proteome that partially overlaps with exercise.

## Introduction

Physical exercise induces coordinated systemic responses that extend beyond contracting skeletal muscle. Acute exercise challenges energy metabolism, vascular regulation, endocrine signaling, immune cell trafficking, and tissue repair. Repeated exposure to these acute perturbations contributes to long-term adaptations in cardiovascular, metabolic, and musculoskeletal physiology. However, the circulating signals that coordinate these responses across tissues remain incompletely understood.

Extracellular vesicles (EVs) are lipid bilayer-delimited particles released by cells and detectable in biofluids such as plasma and serum. EVs contain proteins, lipids, and nucleic acids that can reflect cellular source and participate in intercellular communication (Akers et al., 2013; Whitham et al., 2018). Exercise has emerged as a physiological stimulus capable of increasing circulating EVs and modifying EV cargo, suggesting that EVs may contribute to exercise-induced tissue crosstalk. Whitham et al. provided early evidence that acute exercise promotes the release of EVs containing proteins with potential signaling functions across tissues (Whitham et al., 2018). More recent human studies have shown that the circulating EV proteome responds to different forms of acute exercise. Chong et al. reported changes in a small subset of plasma small-EV proteins following continuous endurance exercise, with responses influenced by aerobic fitness and age (Chong et al., 2024). In contrast, Kobayashi et al. demonstrated rapid changes in the circulating EV proteome after a brief high-intensity interval exercise bout, including proteins associated with coagulation, oxidative stress, and acid–base regulation (Kobayashi et al., 2021). Together, these studies indicate that acute EV protein responses can be detected across distinct exercise modalities, although the influence of long-term training background on these responses remains unclear.

Cold exposure represents another physiological stimulus with marked effects on systemic metabolism and tissue communication. Recent work comparing exercise-trained and cold-exposed inguinal white adipose tissue demonstrated that the two interventions induce distinct proteomic and secretory adaptations, with exercise preferentially increasing proteins related to the extracellular space and vesicle transport, whereas cold exposure produced broader remodeling of the adipose tissue proteome and secretome (Vamvini et al., 2024). These findings suggest that exercise and cold exposure may engage partly distinct, but potentially overlapping, mechanisms of inter-tissue communication. However, whether these stimuli induce shared changes in the circulating EV-enriched proteome has not yet been established.

A key unresolved question is whether acute exercise-induced changes in the circulating EV-enriched proteome are primarily related to the immediate physiological stimulus or are shaped by the individual’s training background. It is also unclear whether acute exercise and cold exposure induce overlapping EV-enriched protein responses in humans. In this exploratory study, we compared serum EV-enriched proteomic profiles before and after acute exercise in endurance-trained athletes, sprinters, natural bodybuilders, and sedentary controls, and related these responses to an independent cold-water immersion dataset. We assessed group-specific protein changes, directionally shared responses, pathway enrichment, and exercise–cold overlap. Our findings indicate that acute EV-enriched proteome remodeling differs across training backgrounds and that cold exposure induces a distinct response with partial directional overlap with exercise.

## Results

### EV proteome profile after acute exercise is dependent on fitness status

To determine whether the training background influences the EV proteome profile, we first isolated the serum of sedentary controls (n = 4), endurance athletes (n = 5), bodybuilders (n = 3) and sprinters (n = 5) before and after a bout of exercise to exhaustion. Small extracellular vesicles were isolated for proteomics as described in Tischer et al., 2025 (Suppl. data 1; Suppl. data 2) (Tischer et al., 2025). Next, group-specific responses were visualized in volcano plots and summarized by the number of significantly increased and decreased proteins (nominal p-value < 0.05) (Fig. 1A-E). From a total of 1395 proteins, 22 proteins were up- and 31 down-regulated in the sedentary group; 27 up- and 21 down-regulated in the endurance athletes group; 22 up- and 25 down-regulated in the bodybuilders group; and 65 up- and 64 down-regulated in the sprinters group (Fig. 1A). Our results show that acute exercise altered the EV proteome in all four training groups. However, the magnitude of the response differed substantially. Sprinters displayed the largest number of significantly altered EV proteins, whereas endurance athletes, bodybuilders and untrained controls showed less changes at the protein level. This supports the idea that the acute EV proteome response is shaped by the training status in humans.

**Figure 1.**
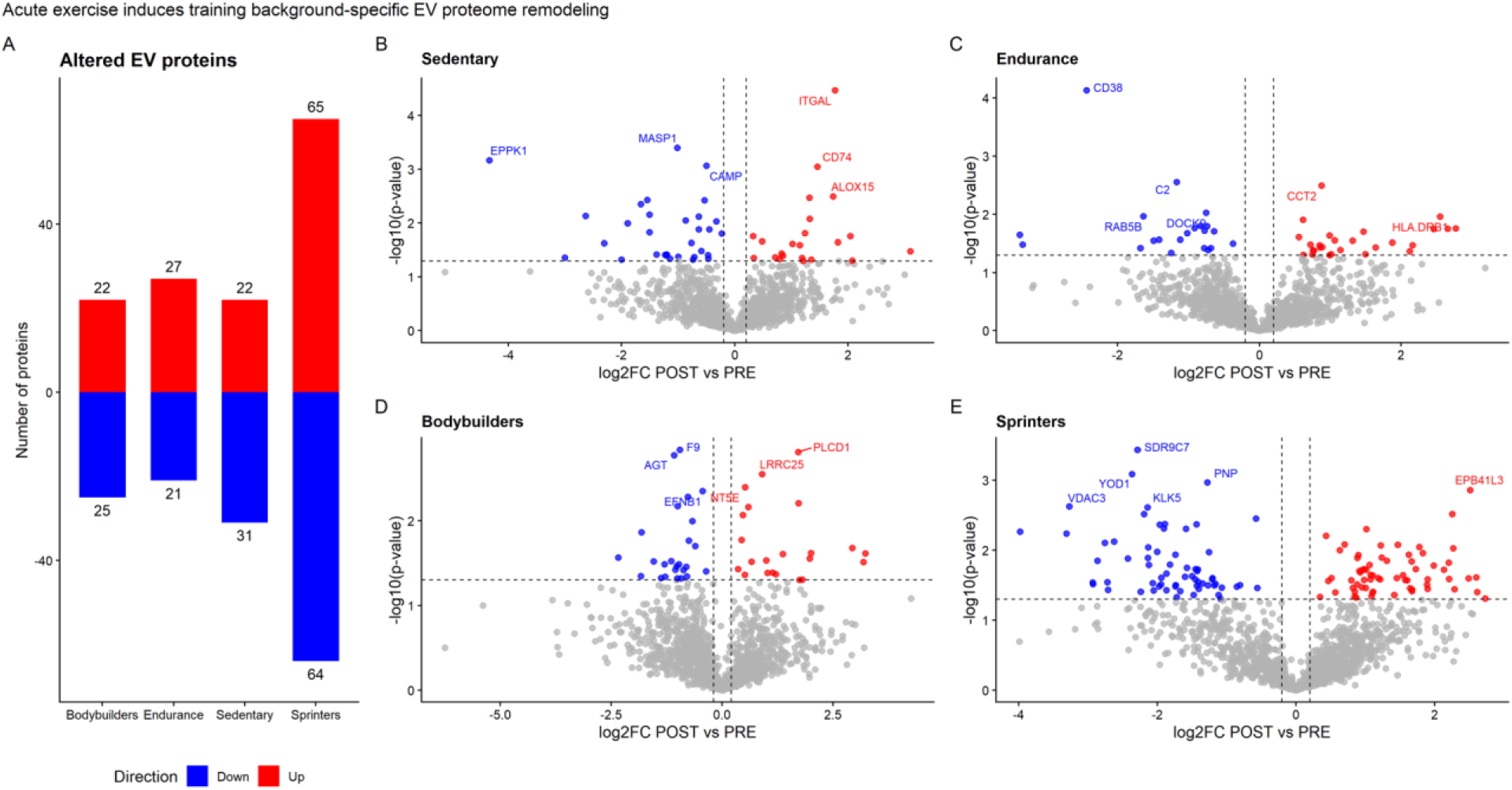
Distinct EV proteome profiles are dependent on fitness status after acute exercise. Diverging bar chart highlighting the different response magnitudes across the four groups (A). Volcano plots of significantly up (red) and down-regulated (blue) serum-derived EV proteins after acute exercise in sedentary controls (B), endurance athletes (C), bodybuilders (D) and sprinters (E). Significance is stated as nominal p-value < 0.05.

### Shared directional EV protein trends across exercise groups

To evaluate whether a common exercise EV protein response existed across training backgrounds, overlap and directionality of significantly regulated proteins between sedentary controls, bodybuilders, endurance athletes and sprinters were analyzed using paired comparisons between post-exercise (POST) versus pre-exercise (PRE) (nominal p-value < 0.05) (Suppl. data 3). Our results show that there is no overlap of nominally significant regulated EV proteins, indicating that no dominant protein signature was statistically conserved across all exercise groups. Because overlap based on statistical significance can be limited in small human intervention studies, we next compared the proteomic response across training groups using a broader approach. To assess shared exercise-associated patterns, we analyzed both significantly and non-significantly regulated proteins showing consistent directionality across all four groups (nominal p-value < 0.05). This allowed us to identify proteins that may represent common exercise-related EV protein trends. Our directionality-based analysis identified proteins that moved in the same direction after exercise in all four groups, even when they may not be individually significant in each group (Fig. 2A). Among a total of 184 proteins that show consistent directionally in the four groups, RGS19, HLA-DRA, CD101, HLA-DRB1, CAND1, EPB41L2, ITGAL and COL4A2 were the most consistently downregulated, whileRAB33B, COLEC11, HRNR, IGHV1.45, ENO3 and SEMG1 were the most consistently upregulated across all training backgrounds (Fig. 2B, 2C).Gene ontology enrichment analysis show that most of these proteins were involved in vesicle trafficking and extracellular matrix organization (Suppl. figure 1; Suppl. data 4).

**Figure 2.**
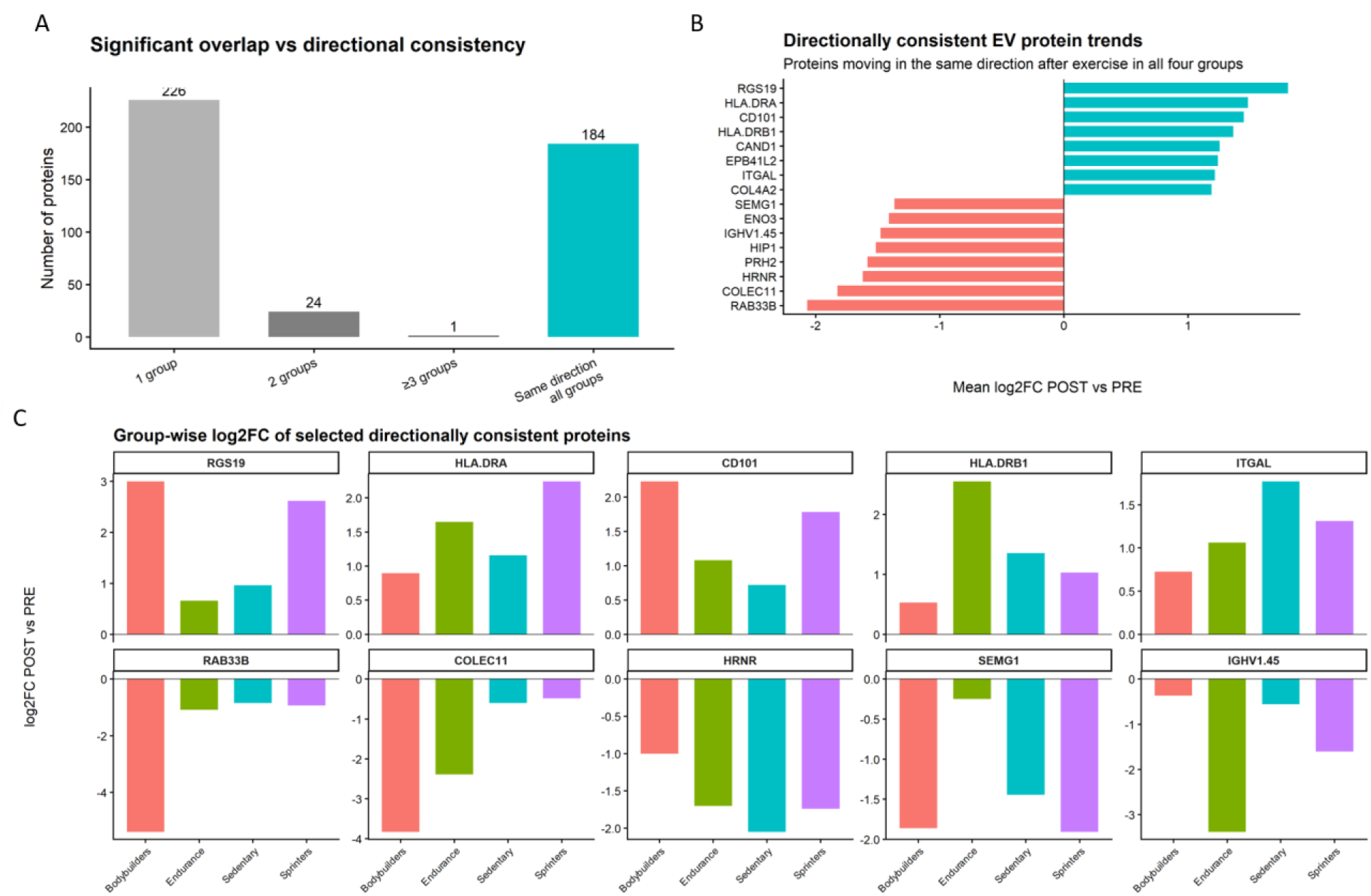
Shared directional EV protein trends across exercise groups. Bar chart showing the number of EV proteins significantly regulated after acute exercise exclusively in one group, 2 groups or ≥ 3 groups and consistently regulated in the same direction in sedentary controls, bodybuilders, endurance athletes and sprinters (A). Top EV proteins showing consistent up-(red) or down (blue) directionality after exercise among the four groups (B). Log_2_ fold change group-wise comparison of selected directionally consistent proteins (C). Significance is stated as nominal p-value < 0.05.

### Cold exposure remodels the EV-enriched proteome and partially overlaps with acute exercise in endurance athletes

To determine whether an acute thermogenic stimulus remodels the circulating EV-enriched proteome, we analyzed serum EVs collected before cold exposure (PRE), immediately after cold exposure (POST), and 30 min after cold exposure (POST30) (n = 8) (Suppl. figure 2). Serum EV-enriched proteomics data from the cold exposure experiment were analyzed using paired comparisons between PRE, POST, and POST30 samples. Proteins meeting a 70% completeness criterion in at least one condition were retained, remaining missing values were imputed using a Perseus-style approach, and paired t-tests were performed for each contrast. Proteins were classified as increased or decreased based on nominal p < 0.05 and the direction of the Log_2_ fold change. FDR-adjusted p-values are reported in the supplementary tables (Suppl. data 5).

Our analysis shows that cold exposure induced time-dependent changes in protein abundance. Immediately after exposure (POST), 76 proteins were significantly altered relative to PRE, including 54 increased and 22 decreased proteins. The POST30 vs POST comparison identified 112 significantly altered proteins, with 28 increased and 84 decreased proteins, while the POST30 vs PRE contrast identified 187 proteins, including 56 increased and 131 decreased proteins (Fig. 4A; Suppl. figures 3-4). Because blood samples from the exercise experiments were also collected immediately after exercise, the cold POST vs PRE contrast was selected for the primary exercise–cold comparison, whereas the POST30 comparisons are presented in the supplementary analyses. Volcano plot analysis of the immediate cold response showed both increased and decreased proteins within the serum EV-enriched fraction (Fig. 4B). Among the nominally increased proteins were IGHG3, LCN2, CP, SERPINA1, and PLVAP, whereas HNRNPA2B1, MCAM, HAGH, DDAH2, and CTNNB1 were among the nominally decreased proteins. These findings indicate that detectable remodeling of the EV proteome was already present immediately after cold exposure. However, the changes were identified using nominal p < 0.05 and should therefore be interpreted as exploratory rather than as FDR-confirmed responses.

It was previously shown that cold exposure acclimatation induces similar physiological effects in the skeletal muscle to endurance exercise in murine models (Johannsen et al., 2026). In this study, we wanted to investigate whether acute cold exposure and acute exercise share similar EV proteomics signatures, particularly within the endurance athletes group. Proteins detected in both datasets were classified according to the concordance of their Log_2_ fold-change direction, without requiring statistical significance in both contrasts. The comparison revealed partially overlapping directional protein trends between endurance exercise and immediate cold exposure (Fig. 4C). Selected proteins increasing in both conditions included KRT82, DYNC1H1, CD247, GP6, LCN2, and RNH1; whereas CD38, RAP2A, CDH1, DOCK9, ZYX, and NAPRT decreased in both conditions (Fig. 4D). None of these selected proteins was nominally significant in both contrasts; rather, each showed a concordant direction of change, with nominal significance in at least one dataset. The overlap therefore represents directionally shared EV-enriched protein trends rather than statistically shared responses. Overall, these exploratory findings suggest that cold exposure and endurance exercise induce partially overlapping acute remodeling of the serum EV-enriched proteome. Equivalent comparisons with sedentary participants, bodybuilders, and sprinters, together with analyses of the POST30 cold-exposure response, are presented in the supplementary material (Suppl. figure 5-6; Suppl. data 6).

**Figure 3.**
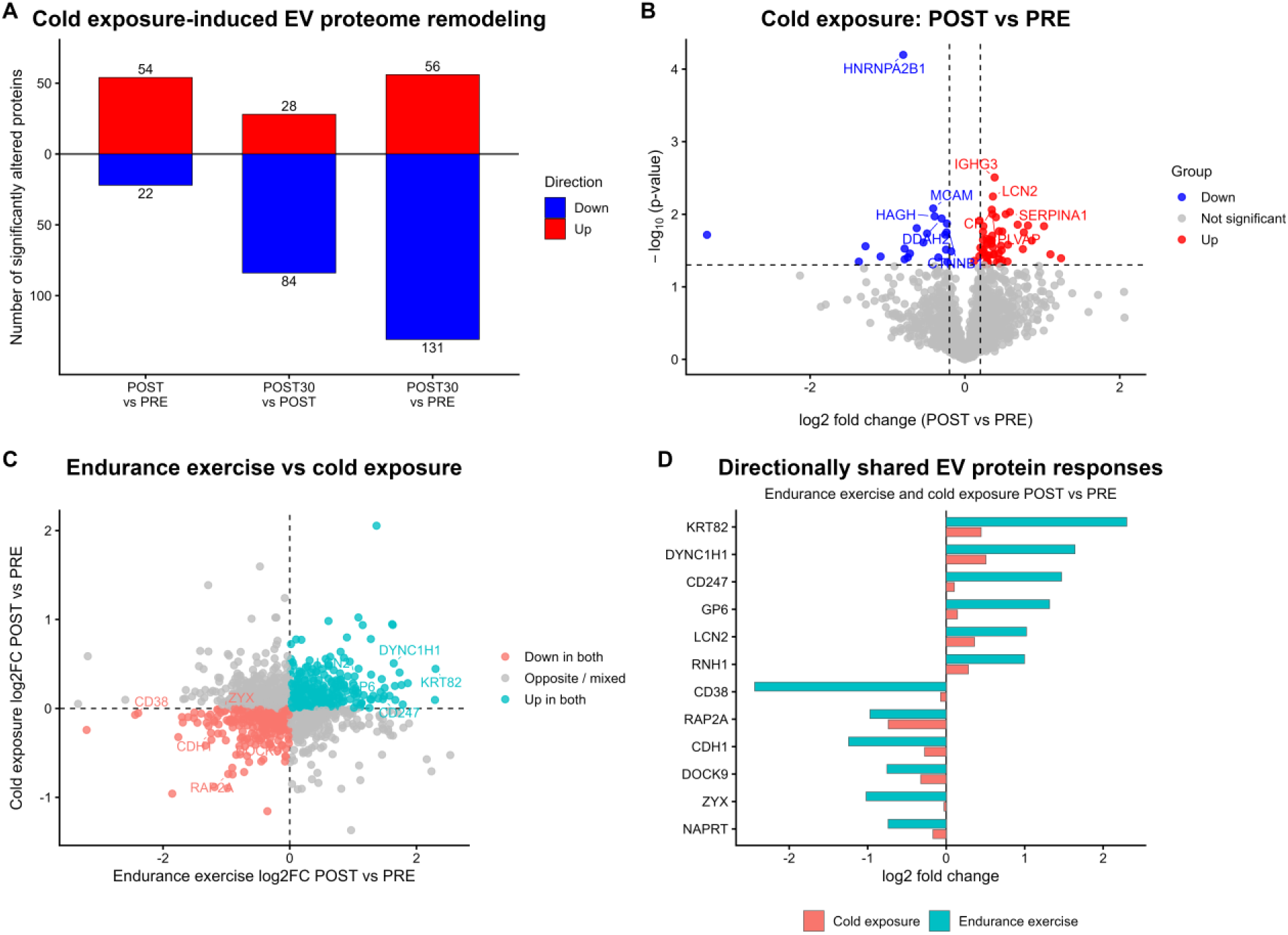
Cold exposure induces time-dependent remodeling of the serum EV proteome and partially overlaps with endurance exercise-induced protein responses. Number of nominally altered proteins in the serum EV for the cold exposure POST vs PRE, POST30 vs POST, and POST30 vs PRE comparisons. Proteins with nominal p < 0.05 and positive Log_2_ fold change are shown as increased, whereas proteins with nominal p < 0.05 and negative Log_2_ fold change are shown as decreased (A). Volcano plot of protein changes immediately after cold exposure compared with PRE. Red indicates increased proteins, blue indicates decreased proteins, and grey indicates proteins that were not nominally significant. Selected proteins are labelled (B). Scatter plot comparing protein changes directly after endurance exercise with changes directly after cold exposure. Proteins increasing in both contrasts are shown in turquoise, proteins decreasing in both are shown in salmon, and proteins with opposite or mixed directions are shown in grey. Selected directionally shared proteins are labelled (C). Selected directionally shared protein responses in POST vs PRE acute exercise in endurance athletes and POST vs PRE cold exposure. Cold exposure Log_2_ fold changes are shown in salmon and endurance exercise Log_2_ fold changes in turquoise. Directional overlap was defined by Log_2_ fold-change signs and did not necessarily indicate statistical significance in both contrasts. All differential comparisons were based on paired tests, with nominal p < 0.05 used for exploratory classification; FDR-adjusted p-values are provided in the supplementary tables (D).

## Discussion

In this exploratory study, we investigated whether acute physiological stress remodels the serum EV-enriched proteome in humans. We compared the proteomic response to acute exercise across individuals with distinct training backgrounds and related these findings to an independent cold exposure dataset. The main observation was that acute exercise induced training-background-specific remodeling of the circulating EV-enriched proteome. Although all exercise groups performed the same acute exercise test, sedentary controls, endurance-trained athletes, bodybuilders, and sprinters showed distinct EV proteomic response patterns. These findings suggest that the circulating EV-enriched proteome is shaped not only by the acute stimulus itself, but also by the long-term physiological and metabolic phenotype of the individual.

The group-specific response patterns support the idea that training background influences systemic communication after exercise. Endurance training, sprint training, resistance-oriented training, and sedentary status are associated with distinct adaptations in skeletal muscle metabolism, vascular function, immune regulation, substrate handling, and endocrine signaling. Circulating EVs have been proposed as mediators of exercise-induced inter-tissue communication, as they can transport proteins, nucleic acids, lipids, and surface molecules between cells and tissues (Nederveen et al., 2021; Whitham et al., 2018). The EV-enriched proteome may therefore reflect differences in tissue stress, cellular activation, EV release, cargo sorting or particle composition between these groups. Importantly, the limited overlap of proteins that reached nominal significance across all exercise groups indicates that there was no single universal EV protein response to acute exercise under these conditions. However, several proteins showed consistent directional changes across groups, suggesting that shared exercise-related EV protein trends may exist even when individual proteins do not reach statistical nominal significance in every group. This distinction is important in small human intervention studies, where strict significance-based overlap may underestimate common biological patterns.

Several exercise-regulated proteins were linked to vesicle trafficking, endosomal biology, immune regulation, complement activation, extracellular matrix organization, cytoskeletal remodeling and stress responses. These categories are biologically plausible in the context of acute exercise, which imposes mechanical, metabolic, vascular and inflammatory stress across multiple tissues. Proteins involved in vesicle trafficking, including Rab proteins, may reflect changes in EV biogenesis, cargo sorting or uptake-related pathways. Rab GTPases are central regulators of intracellular vesicle transport, endosome maturation and membrane trafficking, and are therefore directly relevant to EV biology (Akers et al., 2013; Nagano et al., 2019; Xie et al., 2022). Although the present study cannot determine whether such proteins are functionally active cargo or markers of altered EV origin, their regulation suggests that acute exercise may influence the biology of the circulating EV-enriched particle pool.

The regulation of immune-and complement-associated proteins further supports the concept that acute exercise alters EV-associated systemic signaling. Proteins such as complement components, HLA-related proteins and other immune-associated cargo may reflect transient immune activation, vascular interaction, tissue repair or remodeling processes after exercise. EVs are well-established mediators of immune communication and antigen presentation (Raposo et al., 1996; Robbins & Morelli, 2014), and exercise has previously been shown to alter EV-associated proteins involved in systemic signaling and immune regulation (Maggio et al., 2023; Whitham et al., 2018). In the context of acute exercise, these immune-associated changes should not necessarily be interpreted as pathological inflammation. Instead, they may represent part of the normal acute adaptive response to exercise. This is relevant because metabolic tissues, including adipose tissue and skeletal muscle, are closely linked to immune status during metabolic adaptation and dysfunction (Kawai et al., 2021).

Among the regulated proteins, CD38 was notable because it decreased after exercise in endurance-derived EVs. CD38 is involved in NAD metabolism, immune regulation, and metabolic disease-associated pathways, and has been linked to age-and obesity-related metabolic dysfunction (Benzi, Grozio, et al., 2021; Benzi, Sturla, et al., 2021; Jokinen et al., 2017). Although the functional relevance of EV-associated CD38 was not directly tested in the present study, its regulation may reflect altered NAD-related or inflammatory signaling within the circulating EV-enriched particle pool. This observation should therefore be interpreted as a hypothesis-generating finding rather than evidence for a direct mechanistic role.

To place the exercise findings into a broader physiological context, we also analyzed EV-enriched proteomic changes after cold exposure. Cold exposure represents a distinct acute systemic stressor involving thermoregulatory, sympathetic, vascular and metabolic responses. Similar to exercise, cold exposure induced remodeling of the serum EV-enriched proteome, with dynamic changes across the investigated time points. The comparison between cold exposure and exercise revealed partial overlap in EV-enriched protein responses, suggesting that different acute physiological stressors may share some circulating EV-associated signatures while also inducing stimulus-specific remodeling. This supports the idea that the serum EV-enriched proteome may encode information about both general systemic stress and the specific physiological context of the stimulus.

### Limitations

Several limitations should be considered. First, this was an exploratory study with relatively small group sizes, and the findings should be interpreted as hypothesis-generating. Second, serum EVs isolated by ultracentrifugation are heterogeneous and may contain a smaller proportion of co-isolated lipoproteins, soluble plasma proteins, immunoglobulins, complement proteins, protein aggregates, and other extracellular particles. Therefore, the results should be interpreted as changes in the serum EV-enriched particle proteome rather than as a highly pure EV proteome. Third, the study design does not allow the cellular or tissue origin of individual proteins to be assigned with certainty. Future studies combining EV subpopulation separation, cell-type-specific markers, orthogonal EV characterization, and larger cohorts would be required to determine whether the observed protein changes reflect altered EV release, altered cargo sorting or changes in the composition of circulating particle populations. Finally, the cold exposure dataset was used as an independent physiological comparison rather than a matched validation cohort, and differences in stimulus type, sampling time, and participant characteristics limit direct interpretation. Given the exploratory nature of the study, the limited sample size, and the expected variability of serum EV proteomics, nominally significant proteins were interpreted cautiously.

Protein-level findings were therefore considered hypothesis-generating and were supported by directional trends, group-wise comparisons, and pathway enrichment analyses rather than treated as definitive mechanistic endpoints. In addition, the fact that each training group present a different set of significantly EV proteins after exercise when analysed separately does not necessarily imply a difference between groups.

Taken together, this study shows that acute exercise and cold exposure remodel the serum EV-enriched proteome in humans. The exercise response was strongly influenced by training background, indicating that long-term physiological adaptation shapes the circulating EV-enriched proteomic response to an acute stimulus. The partial overlap between exercise and cold exposure further suggests that circulating EV-enriched particles may reflect both shared acute-stress responses and stimulus-specific systemic remodeling. These findings provide an exploratory resource for future studies investigating how exercise training status and acute physiological stress influence EV-associated inter-tissue communication in humans.

### Conclusion

Acute exercise induces remodeling of the serum EV-enriched proteome in humans, but the response is not uniform across training backgrounds. Sprinters showed the largest acute response, while endurance athletes, bodybuilders and untrained controls displayed distinct proteomic patterns. Although few proteins were significantly shared across all groups, a broader set of proteins showed concordant directional trends, suggesting a modest shared EV protein response superimposed on training-specific remodeling. Cold exposure induced an additional time-dependent EV proteome response with partial but incomplete overlap with exercise. Together, these findings support the use of circulating EV-enriched proteomics as an exploratory readout of acute physiological stress and training-state-dependent systemic adaptation.

## Methods

### Study design and participants

This exploratory study analyzed serum EV-enriched proteomes from male participants representing four training backgrounds: untrained sedentary controls (n = 4), endurance athletes (n = 5), natural bodybuilders (n = 3), and sprinters (n = 5). Participants underwent baseline blood sampling followed by a maximal bicycle ramp test and post-exercise blood sampling (Schranner et al., 2021). Physiological characterization including parameters such as age, body weight, BMI, VO2max or VO2M, ventilatory threshold, leg strength, relative leg strength, grip strength, and reactive strength can be found in Schranner et al. and in the table below (Schranner et al., 2021):

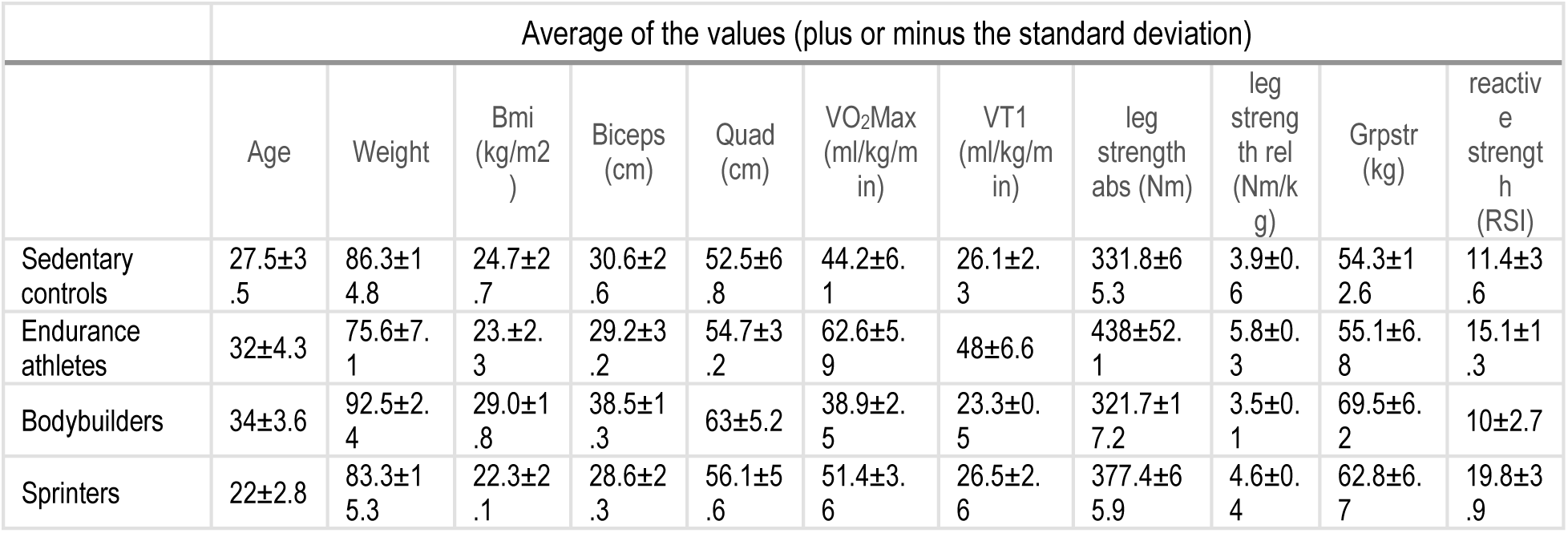

|  | Average of the values (plus or minus the standard deviation) |  |  |  |  |  |  |  |  |  |
|---|---|---|---|---|---|---|---|---|---|---|
|  | Age | Weight | Bmi (kg/m2) | Biceps (cm) | Quad (cm) | VO<sub>2</sub>Max (ml/kg/min) | VT1 (ml/kg/min) | leg strength abs (Nm) | leg strength rel (Nm/kg) | Grpstr (kg) | reactive strength (RSI) |
| Sedentary controls | 27.5±3.5 | 86.3±14.8 | 24.7±2.7 | 30.6±2.6 | 52.5±6.8 | 44.2±6.1 | 26.1±2.3 | 331.8±65.3 | 3.9±0.6 | 54.3±12.6 | 11.4±3.6 |
| Endurance athletes | 32±4.3 | 75.6±7.1 | 23.±2.3 | 29.2±3.2 | 54.7±3.2 | 62.6±5.9 | 48±6.6 | 438±52.1 | 5.8±0.3 | 55.1±6.8 | 15.1±1.3 |
| Bodybuilders | 34±3.6 | 92.5±2.4 | 29.0±1.8 | 38.5±1.3 | 63±5.2 | 38.9±2.5 | 23.3±0.5 | 321.7±17.2 | 3.5±0.1 | 69.5±6.2 | 10±2.7 |
| Sprinters | 22±2.8 | 83.3±15.3 | 22.3±2.1 | 28.6±2.3 | 56.1±5.6 | 51.4±3.6 | 26.5±2.6 | 377.4±65.9 | 4.6±0.4 | 62.8±6.7 | 19.8±3.9 |

Cold exposure data was derived from an independent ice bath study in which participants (n = 8, 3 males and 5 females; age average: 24.9 ± 4.9 years old; BMI: 22.8 ± 1.7 kg/m^2^) underwent cold-water immersion at 8-12°C for 5 min. Details about the human cold exposure intervention can be found in Braunsperger et al., 2025 (Braunsperger et al., 2025). Blood samples were collected before, immediately after, and 30 min after cold exposure (Braunsperger et al., 2025). This dataset was used as an independent acute physiological stress comparison rather than as a direct mechanistic validation cohort.

### Blood collection and serum preparation

For the exercise and cold exposure cohort, blood was allowed to clot at room temperature for 10 min and centrifuged at 2,460 g for 10 minutes at 18°C. Serum was aliquoted and stored at -80 until EV-enriched particle isolation.

### EV isolation by ultracentrifugation

For the proteomics analyses serum EVs were isolated by ultracentrifugation. Briefly, serum was centrifuged at 2,460 g for 10 minutes at 15°C to remove cells and debris. Next, samples were centrifuged at 20,000 g for 40 minutes at 4°C, supernatant was collected and small EVs were pellet by high-speed ultracentrifugation at 100,000 g for 2 hours 4°C. Pellets were washed in PBS and re-centrifuged at 100,000 g for 2 hours 4°C before being resuspended in PBS for EV characterization and proteomics sample preparation.

### EV characterization by nanoparticle tracking analysis and electron microscopy

ZetaView nanoparticle tracking analyzer (Particle Metrix, Germany) was used for concentration and size distribution of small EVs isolated by ultracentrifugation. Transmission electron microscopy was used for the visualization of small EVs.

For electron microscopy, samples were bound to glow discharged formvar/carbon-coated copper TEM grids (400 mesh) (Science Services, München, Germany). Loaded TEM grids were then incubated on drops of 1% glutaraldehyde (Science Services, München, Germany) for fixation followed by eight times washing on drops of filtered ultrapure distilled water (Ampuwa, Fresenius Kabi, Bad Homburg, Germany). Immediately after washing the TEM grids were negatively stained with 2% uranyl acetate (Science Services, München, Germany) for 45 – 60 seconds and air dried. Images were acquired with a Crossbeam 550 (Carl Zeiss, Oberkochen, Germany) with STEM detector at 30 kV acceleration voltage, 150 pA current in imaging mode.

EV characterization results and TEM images from the exercise intervention are shown in Tischer et al.(Tischer et al., 2025). For the cold exposure study, EVs characterization can be found in Suppl. fig. 2.

### Proteomics sample preparation and LC-MS/MS analysis

Proteomics raw data of exercise-induced EVs were processed as described in Tischer et al. (Tischer et al., 2025). EV-enriched protein samples were processed for LC-MS/MS-based proteomics by lysis in urea lysis buffer (4 M urea, 10 mM Bond-Breaker TCEP solution [Thermo], 30 mM CAA [Merck] in Tris-HCl pH 8.0). After incubation at room temperature and 800 rpm on a thermo shaker for 30 minutes, proteins were enriched, desalted, and prepared for enzymatic digestion using the single-pot solid-phase-enhanced sample preparation (SP3) protocol (Hughes et al., 2019). Samples were incubated with 2 µl of SP3 magnetic beads and 100 µl of acetonitrile (ACN, [VWR]) on a thermo shaker at 1,000 rpm and room temperature for 5 min. Following, protein-bead aggregates were captured on a magnetic rack, the supernatant was removed and discarded, and beads were washed three times with 180 µl of 80 % ethanol. The beads were briefly air-dried and resuspended in 10 µl digestion buffer (0.1 µg trypsin/Lys-C [Promega] in 50 mM TEAB). The enzymatic digestion was carried out at 37 °C and 800 rpm in a thermo shaker for 18 hours. The next day, tryptic peptides were desalted by adding 200 µl of pure ACN, followed by incubation at 800 rpm and room temperature for 5 minutes. After bead capturing on a magnetic rack, the supernatants were discarded, and bead-peptide aggregates rinsed with 200 µl ACN. Peptides were eluted from beads with 10 µl of 5 % ACN in LC-MS-grade water [VWR] and acidified by adding 2 µl of 10 % formic acid [Sigma]. After peptide concentration determination using a Nanodrop One system [Thermo, 200 ng of peptides were loaded onto an Evotip Pure [Evosep] trap column following the manufacturer’s protocol.

### LC-MS/SM analysis

Proteomics acquisition was carried out with an Evosep One chromatographic system [Evosep] and a timsTOF Pro 2 mass spectrometer (Bruker Daltonics). Using the preinstalled 40-samples-per-day whisper method, peptides were separated on an Aurora Elite CSI analytical column (length: 15 cm, inner diameter: 75 µm, 1.7 µm C18 particles [Ionopticks] within 31 min. Eluting peptides were transferred into the mass spectrometer via a Captive Spray nano-electrospray ion source operated at a constant spray voltage of 1.5 kV. MS measurements were carried out in DIA-PASEF mode (data-independent acquisition parallel accumulation-serial fragmentation). Ion mobility-based separation was carried out in the dual TIMS analyser operating with 100 ms accumulation and ramping times and a 100% duty cycle. Selected peptide ion species were restricted to a mass-to-charge range of 400 – 1000 m/z and an ion mobility range of 0.70 – 1.35 Vs/cm2. The DIA isolation space consisted of 24 equal-sized PASEF windows with a width of 25 m/z each. For fragmentation, collisional energies were stepwise increased from 24 eV at 0.7 Vs/cm2 to 49 eV at 1.35 Vs/cm2.

The mass spectrometry proteomics data have been deposited to the ProteomeXchange Consortium (http://proteomecentral.proteomexchange.org) via the PRIDE partner repository (Perez-Riverol et al., 2022) with the dataset identifiers **PXD058619** (EV proteomics before and after exercise) and **PXD087502** (EV proteomics before and after cold exposure).

### Differential protein abundance analysis

Differential protein abundance was analyzed separately for each training group using paired comparisons of post-exercise versus baseline EV-enriched protein abundance. For the cold exposure dataset, paired analyses were performed for three contrasts: POST vs PRE, POST30 vs POST, and POST30 vs PRE. For each contrast, Log_2_ fold changes were calculated as the mean paired difference between conditions, and statistical significance was assessed using paired tests. Proteins were classified as increased when Log_2_FC > 0 with nominal p < 0.05, decreased when Log_2_FC < 0 with nominal p < 0.05, and not significantly altered when p ≥ 0.05. No additional absolute Log_2_FC threshold was applied for the primary classification of altered proteins.

For comparisons between exercise and cold exposure, proteins shared between the respective exercise and cold exposure datasets were matched by protein symbol. Directionally shared protein trends were defined based on concordant log2FC direction between contrasts. Proteins with positive log2FC values in both contrasts were classified as “up in both,” proteins with negative log2FC values in both contrasts as “down in both,” and proteins with opposing directions as “opposite/mixed.” These directional comparisons were used to evaluate overlap in EV protein response patterns and were not restricted to proteins reaching statistical significance in both contrasts. The main exercise–cold comparison focused on endurance exercise versus cold exposure POST30 vs PRE, while analogous comparisons for sedentary participants, bodybuilders, and sprinters were included as supplementary analyses. Multiple testing correction was calculated using the Benjamini–Hochberg procedure and reported where applicable.

### Statistics and reproducibility

All statistical analyses and visualizations were performed in R. Paired tests were used for within subject exercise and cold exposure comparisons where appropriate. Differentially regulated proteins were classified using nominal p < 0.05 and Log2FC direction, without applying an additional absolute Log2FC cutoff. The complete R scripts, package versions, package source links, and session information are provided in the supplementary material (Suppl. data 7).

## CRediT authorship contribution statement

**Dominik Tischer**: Writing – review & editing, Writing – original draft, Visualization, Methodology, Investigation, Formal analysis, Data curation, Conceptualization. **Alexander Braunsperger**: Writing – review & editing, Investigation, Resources (human cold exposure study). **Sebastian Kallabis**: Writing – review & editing, Methodology, Investigation (proteomics). **Daniela Schranner**: Resources (human exercise study). **Paul Jonas Jost**: Investigation. **Jan Hasenauer**: Supervision. **Felix Meissner**: Supervision. **Alexander Pfeifer**: Writing – review & editing, Supervision, Resources, Funding acquisition. **Henning Wackerhage**: Writing – review & editing, Supervision, Resources, Funding acquisition. **Ana Soriano-Arroquia**: Writing – review & editing, Supervision, Project administration, Methodology, Investigation, Funding acquisition, Conceptualization.

## Acknowledgments

This work was funded by the Deutsche Forschungsgemeinschaft (DFG, German Research Foundation) – TRR333/1 – project number 450149205 Transregio collaborative research center BATenergy to A.S.-A. and H.W. D.S was supported with a doctoral scholarship of the German National Scholarship Foundation (Studienstiftung des Deutschen Volkes). We thank Timo Wadenpohl and Prof. Dr. Stephanie Jung for the use of the Zetaview system and the Microscopy Core Facility of the Medical Faculty at the University of Bonn for providing support and instrumentation funded by the Deutsche Forschungsgemeinschaft (DFG, German Research Foundation) – project number 388171357. Statistical analyses were performed using R version 4.5.3 and GraphPad Prism version 6 for Windows, GraphPad Software, Boston, Massachusetts USA, www.graphpad.com.

