## Supplementary material for "Training background shapes the exercise-induced extracellular vesicle proteome to acute exercise with partial overlap with cold exposure in humans": Suppl_Data_7_R__scripts.pdf

### Complete R Analysis Script

#### Serum EV-enriched proteomics: exercise and cold exposure

**Important scope note.** The exercise part uses the processed and normalized group-specific differential-result tables. The cold-exposure part is re-derived from cold_raw.xlsx using one common 70%-filtered and imputed matrix. Nominal $p < 0.05$ is used for the exploratory figure classification, while Benjamini-Hochberg FDR values are calculated and saved in all result tables.

#### Required input files

- cold_raw.xlsx
- Sedentary_EV_Proteomics_Results.csv
- Endurance_EV_Proteomics_Results.csv
- Bodybuilders_EV_Proteomics_Results.csv
- Sprinters_EV_Proteomics_Results.csv

**Software:** R with the packages readxl, dplyr, tidyr, ggplot2, ggrepel, patchwork, writexl, and gprofiler2.

**Recommended use:** Save the code below as one .R file in the folder containing the input files, then run from top to bottom in a clean R session.

#### Analysis sections

1. Package setup and helper functions

2. Exercise group-specific POST vs PRE analyses

3. Cold-exposure common-matrix filtering, imputation, and paired tests

4. Cold PCA analyses

5. Cold volcano plots

6. Exercise-cold directional comparisons

7. Main manuscript Figure 4

8. Thesis endurance-cold figures

9. Supplementary GO Biological Process analysis

10. Consolidated workbook and reproducibility outputs

#### Complete R script

**Copy the entire code block without the line numbers.** The script creates a Final_EV_Proteomics_Analysis folder containing all result tables, figures, a consolidated Excel workbook, session information, and an R workspace.

```
# =============================================================================
# COMPLETE R ANALYSIS SCRIPT
# Serum EV-enriched proteomics: exercise groups and cold exposure
# =============================================================================
#
# Analysis scope
# --------------
# 1. Exercise analyses use the previously processed and normalized group-specific
#    differential-abundance result tables supplied by the proteomics facility.
#    Raw exercise MS data are not reprocessed in this script.
# 2. Cold-exposure differential abundance is re-derived from cold_raw.xlsx so
#    POST vs PRE, POST30 vs PRE, and POST30 vs POST use one common protein matrix.
# 3. Primary exploratory classification:
#       Up   = log2FC > 0 and nominal p < 0.05
#       Down = log2FC < 0 and nominal p < 0.05
#       Not significant = p >= 0.05
#    Benjamini-Hochberg FDR values are calculated and retained in all result
#    tables. No additional absolute log2FC threshold is used for classification.
# 4. Exercise-cold directional overlap is based on the sign of log2FC and does
#    not require significance in both contrasts.
# 5. Terminology: serum EV-enriched fractions / EV-enriched particles.
#
# Expected input files
# --------------------
# cold_raw.xlsx
# Sedentary_EV_Proteomics_Results.csv
# Endurance_EV_Proteomics_Results.csv
# Bodybuilders_EV_Proteomics_Results.csv
# Sprinters_EV_Proteomics_Results.csv
#
# Output folders are created automatically.
# =============================================================================

# -----------------------------------------------------------------------------
# 0. Packages and reproducibility settings
# -----------------------------------------------------------------------------

required_packages <- c(
  "readxl", "dplyr", "tidyr", "ggplot2", "ggrepel", "patchwork",
  "writexl", "gprofiler2"
)

missing_packages <- required_packages[
  !vapply(required_packages, requireNamespace, logical(1), quietly = TRUE)
]

if (length(missing_packages) > 0) {
  install.packages(missing_packages)
}

library(readxl)
library(dplyr)
library(tidyr)
library(ggplot2)
library(ggrepel)
library(patchwork)
library(writexl)
library(gprofiler2)

set.seed(123)

output_dir <- "Final_EV_Proteomics_Analysis"
dir.create(output_dir, recursive = TRUE, showWarnings = FALSE)
dir.create(file.path(output_dir, "tables"), recursive = TRUE, showWarnings = FALSE)
dir.create(file.path(output_dir, "figures"), recursive = TRUE, showWarnings = FALSE)
dir.create(file.path(output_dir, "supplementary_GO"), recursive = TRUE, showWarnings = FALSE)

# -----------------------------------------------------------------------------
# 1. General helper functions
# -----------------------------------------------------------------------------

classify_change <- function(log2FC, p_value) {
  dplyr::case_when(
    !is.na(p_value) & p_value < 0.05 & log2FC > 0 ~ "Up",
    !is.na(p_value) & p_value < 0.05 & log2FC < 0 ~ "Down",
```

```
    TRUE ~ "Not significant"
  )
}

standardize_exercise_results <- function(file, group_name) {
  x <- read.csv(file, check.names = FALSE, stringsAsFactors = FALSE)

  required <- c("Protein", "log2FC", "p_value")
  if (!all(required %in% names(x))) {
    stop("Missing required columns in ", file, ": ",
         paste(setdiff(required, names(x)), collapse = ", "))
  }

  x %>%
    transmute(
      Protein = trimws(as.character(Protein)),
      log2FC = suppressWarnings(as.numeric(log2FC)),
      p_value = suppressWarnings(as.numeric(p_value)),
      FDR = if ("FDR" %in% names(x)) suppressWarnings(as.numeric(FDR))
            else p.adjust(suppressWarnings(as.numeric(p_value)), method = "BH"),
      Group = group_name
    ) %>%
    filter(
      !is.na(Protein), Protein != "",
      !is.na(log2FC), !is.na(p_value)
    ) %>%
    distinct(Protein, .keep_all = TRUE) %>%
    mutate(
      Regulation_nominal = classify_change(log2FC, p_value),
      Regulation_FDR = classify_change(log2FC, FDR),
      neg_log10_p = -log10(pmax(p_value, .Machine$double.xmin))
    )
}

impute_sample_perseus <- function(x, downshift = 1.8, width = 0.3) {
  missing_idx <- which(is.na(x))
  if (length(missing_idx) == 0) return(x)

  observed <- x[!is.na(x)]
  if (length(observed) < 2) return(x)

  mu <- mean(observed)
  sigma <- sd(observed)
  if (!is.finite(sigma) || sigma == 0) sigma <- 0.01

  x[missing_idx] <- rnorm(
    length(missing_idx),
    mean = mu - downshift * sigma,
    sd = width * sigma
  )
  x
}

paired_protein_test <- function(matrix_data, metadata, condition_2, condition_1) {
  subjects <- intersect(
    as.character(metadata$Subject[metadata$Timepoint == condition_1]),
    as.character(metadata$Subject[metadata$Timepoint == condition_2])
  )

  subjects <- sort(unique(subjects))

  get_condition_matrix <- function(tp) {
    idx <- metadata$Timepoint == tp & as.character(metadata$Subject) %in% subjects
    m <- as.matrix(matrix_data[idx, , drop = FALSE])
    rownames(m) <- as.character(metadata$Subject[idx])
    m[subjects, , drop = FALSE]
  }

  m1 <- get_condition_matrix(condition_1)
  m2 <- get_condition_matrix(condition_2)

  log2FC <- colMeans(m2 - m1, na.rm = TRUE)

  p_value <- vapply(seq_len(ncol(m1)), function(j) {
    x <- m2[, j]
    y <- m1[, j]
    keep <- is.finite(x) & is.finite(y)
    if (sum(keep) < 2) return(NA_real_)
    tryCatch(t.test(x[keep], y[keep], paired = TRUE)$p.value,
             error = function(e) NA_real_)
  }, numeric(1))
```

```
  n_pairs <- vapply(seq_len(ncol(m1)), function(j) {
    sum(is.finite(m2[, j]) & is.finite(m1[, j]))
  }, integer(1))

  out <- data.frame(
    Protein = colnames(m1),
    Contrast = paste0(condition_2, " vs ", condition_1),
    log2FC = log2FC,
    p_value = p_value,
    n_pairs = n_pairs,
    stringsAsFactors = FALSE
  )

  out$FDR <- p.adjust(out$p_value, method = "BH")
  out$Regulation_nominal <- classify_change(out$log2FC, out$p_value)
  out$Regulation_FDR <- classify_change(out$log2FC, out$FDR)
  out$neg_log10_p <- -log10(pmax(out$p_value, .Machine$double.xmin))
  out
}

# ---------------------------------------------------------------------------
# 2. Exercise group-specific POST vs PRE result tables
# ---------------------------------------------------------------------------

exercise_files <- c(
  Sedentary = "Sedentary_EV_Proteomics_Results.csv",
  Endurance = "Endurance_EV_Proteomics_Results.csv",
  Bodybuilders = "Bodybuilders_EV_Proteomics_Results.csv",
  Sprinters = "Sprinters_EV_Proteomics_Results.csv"
)

exercise_results <- lapply(names(exercise_files), function(group_name) {
  standardize_exercise_results(exercise_files[[group_name]], group_name)
})
names(exercise_results) <- names(exercise_files)

exercise_all <- bind_rows(exercise_results)

exercise_summary <- exercise_all %>%
  count(Group, Regulation_nominal, name = "n") %>%
  tidyr::pivot_wider(
    names_from = Regulation_nominal,
    values_from = n,
    values_fill = 0
  )

write.csv(
  exercise_all,
  file.path(output_dir, "tables", "Exercise_all_groups_standardized_results.csv"),
  row.names = FALSE
)
write.csv(
  exercise_summary,
  file.path(output_dir, "tables", "Exercise_group_nominal_change_summary.csv"),
  row.names = FALSE
)

# Exercise volcano plots
exercise_volcano_plot <- function(data, title_text) {
  ggplot(data, aes(log2FC, neg_log10_p, colour = Regulation_nominal)) +
    geom_point(size = 1.8, alpha = 0.75) +
    geom_hline(yintercept = -log10(0.05), linetype = "dashed", linewidth = 0.4) +
    geom_vline(xintercept = 0, linetype = "dashed", linewidth = 0.4) +
    scale_colour_manual(values = c(
      "Down" = "blue", "Not significant" = "grey70", "Up" = "red"
    )) +
    labs(
      title = title_text,
      x = "log2 fold change (POST vs PRE)",
      y = expression(-log[10](p)),
      colour = NULL
    ) +
    theme_classic(base_size = 12) +
    theme(plot.title = element_text(face = "bold", hjust = 0.5))
}

exercise_volcanoes <- lapply(names(exercise_results), function(g) {
  exercise_volcano_plot(exercise_results[[g]], paste0(g, ": POST vs PRE"))
})
names(exercise_volcanoes) <- names(exercise_results)

exercise_volcano_figure <-
```

```
  (exercise_volcanoes$Sedentary | exercise_volcanoes$Endurance) /
  (exercise_volcanoes$Bodybuilders | exercise_volcanoes$Sprinters) +
  plot_annotation(tag_levels = "A")

ggsave(
  file.path(output_dir, "figures", "Exercise_group_volcanoes.png"),
  exercise_volcano_figure, width = 14, height = 10, dpi = 600, bg = "white"
)
ggsave(
  file.path(output_dir, "figures", "Exercise_group_volcanoes.pdf"),
  exercise_volcano_figure, width = 14, height = 10, bg = "white"
)

# ---------------------------------------------------------------------------
# 3. Cold exposure: common-matrix processing and paired analyses
# ---------------------------------------------------------------------------

cold_raw <- readxl::read_excel("cold_raw.xlsx")

# Required metadata columns
required_meta <- c("Subject", "Timepoint")
if (!all(required_meta %in% names(cold_raw))) {
  stop("cold_raw.xlsx must contain Subject and Timepoint columns.")
}

cold_raw$Subject <- as.character(cold_raw$Subject)
cold_raw$Timepoint <- toupper(trimws(as.character(cold_raw$Timepoint)))

# Keep subjects represented at PRE, POST and POST30
subject_timepoint_table <- table(cold_raw$Subject, cold_raw$Timepoint)
needed_timepoints <- c("PRE", "POST", "POST30")
if (!all(needed_timepoints %in% colnames(subject_timepoint_table))) {
  stop("Expected PRE, POST and POST30 in Timepoint.")
}

complete_subjects <- rownames(subject_timepoint_table)[
  apply(subject_timepoint_table[, needed_timepoints, drop = FALSE], 1,
        function(x) all(x >= 1))
]

cold_raw <- cold_raw[cold_raw$Subject %in% complete_subjects, , drop = FALSE]

metadata_columns <- intersect(
  c("Sample", "Subject", "Sex", "Timepoint"),
  names(cold_raw)
)

cold_meta <- as.data.frame(cold_raw[, metadata_columns, drop = FALSE])
protein_columns <- setdiff(names(cold_raw), metadata_columns)

cold_matrix <- as.data.frame(cold_raw[, protein_columns, drop = FALSE])
cold_matrix[] <- lapply(cold_matrix, function(x) suppressWarnings(as.numeric(x)))

# Remove unnamed, entirely missing and zero-information columns
valid_name <- !is.na(names(cold_matrix)) & names(cold_matrix) != "" &
  !grepl("^\\.\\.\\.|^Unnamed", names(cold_matrix), ignore.case = TRUE)
not_all_missing <- vapply(cold_matrix, function(x) any(is.finite(x)), logical(1))
cold_matrix <- cold_matrix[, valid_name & not_all_missing, drop = FALSE]

# 70% completeness criterion in at least one condition
n_subjects <- length(unique(cold_meta$Subject))
completeness_threshold <- ceiling(0.70 * n_subjects)

valid_counts <- lapply(needed_timepoints, function(tp) {
  colSums(!is.na(cold_matrix[cold_meta$Timepoint == tp, , drop = FALSE]))
})
names(valid_counts) <- needed_timepoints

keep_proteins <-
  valid_counts$PRE >= completeness_threshold |
  valid_counts$POST >= completeness_threshold |
  valid_counts$POST30 >= completeness_threshold

cold_matrix_filtered <- cold_matrix[, keep_proteins, drop = FALSE]

cold_filter_summary <- data.frame(
  Protein = names(cold_matrix),
  valid_PRE = valid_counts$PRE,
  valid_POST = valid_counts$POST,
  valid_POST30 = valid_counts$POST30,
  threshold = completeness_threshold,
  passed_70percent_filter = keep_proteins,
```

```
  stringsAsFactors = FALSE
)

# Sample-wise Perseus-style imputation
cold_matrix_imputed <- t(apply(
  as.matrix(cold_matrix_filtered), 1, impute_sample_perseus,
  downshift = 1.8, width = 0.3
))
cold_matrix_imputed <- as.data.frame(cold_matrix_imputed, check.names = FALSE)

# Remove proteins with zero variance after imputation
protein_sd <- apply(cold_matrix_imputed, 2, sd)
keep_variable <- is.finite(protein_sd) & protein_sd > 0
cold_matrix_imputed <- cold_matrix_imputed[, keep_variable, drop = FALSE]

cold_post_vs_pre <- paired_protein_test(
  cold_matrix_imputed, cold_meta, "POST", "PRE"
)
cold_post30_vs_pre <- paired_protein_test(
  cold_matrix_imputed, cold_meta, "POST30", "PRE"
)
cold_post30_vs_post <- paired_protein_test(
  cold_matrix_imputed, cold_meta, "POST30", "POST"
)

write.csv(
  cold_filter_summary,
  file.path(output_dir, "tables", "Cold_70percent_filtering_summary.csv"),
  row.names = FALSE
)
write.csv(
  cbind(cold_meta, cold_matrix_imputed),
  file.path(output_dir, "tables", "Cold_common_matrix_filtered_imputed.csv"),
  row.names = FALSE
)
write.csv(
  cold_post_vs_pre,
  file.path(output_dir, "tables", "Cold_POST_vs_PRE_same_matrix_results.csv"),
  row.names = FALSE
)
write.csv(
  cold_post30_vs_pre,
  file.path(output_dir, "tables", "Cold_POST30_vs_PRE_same_matrix_results.csv"),
  row.names = FALSE
)
write.csv(
  cold_post30_vs_post,
  file.path(output_dir, "tables", "Cold_POST30_vs_POST_same_matrix_results.csv"),
  row.names = FALSE
)

cold_summary <- bind_rows(
  cold_post_vs_pre %>% count(Contrast, Regulation_nominal, name = "n"),
  cold_post30_vs_pre %>% count(Contrast, Regulation_nominal, name = "n"),
  cold_post30_vs_post %>% count(Contrast, Regulation_nominal, name = "n")
)
write.csv(
  cold_summary,
  file.path(output_dir, "tables", "Cold_timecourse_nominal_change_summary.csv"),
  row.names = FALSE
)

# ----------------------------------------------------------------------------
# 4. Cold PCA analyses
# ----------------------------------------------------------------------------

make_two_timepoint_pca <- function(abundance_matrix, metadata, tp1, tp2, title_text) {
  keep_samples <- metadata$Timepoint %in% c(tp1, tp2)
  x <- abundance_matrix[keep_samples, , drop = FALSE]
  meta <- metadata[keep_samples, , drop = FALSE]

  sds <- apply(x, 2, sd)
  x <- x[, is.finite(sds) & sds > 0, drop = FALSE]

  pca <- prcomp(x, center = TRUE, scale. = TRUE)
  variance <- 100 * pca$sdev^2 / sum(pca$sdev^2)

  scores <- data.frame(
    meta,
    PC1 = pca$x[, 1],
    PC2 = pca$x[, 2],
    stringsAsFactors = FALSE
```

```
  )
  scores$Timepoint <- factor(scores$Timepoint, levels = c(tp1, tp2))

  p <- ggplot(scores, aes(PC1, PC2, colour = Timepoint)) +
    geom_point(size = 4, alpha = 0.85, shape = 16) +
    labs(
      title = title_text,
      x = paste0("PC1 (", round(variance[1], 1), "%)"),
      y = paste0("PC2 (", round(variance[2], 1), "%)"),
      colour = "Timepoint"
    ) +
    theme_classic(base_size = 14) +
    theme(plot.title = element_text(face = "bold", hjust = 0.5))

  list(plot = p, scores = scores, pca = pca, variance = variance)
}

pca_pre_post <- make_two_timepoint_pca(
  cold_matrix_imputed, cold_meta, "PRE", "POST",
  "Cold exposure: PRE vs POST"
)
pca_pre_post30 <- make_two_timepoint_pca(
  cold_matrix_imputed, cold_meta, "PRE", "POST30",
  "Cold exposure: PRE vs POST30"
)

pca_colours <- c("PRE" = "#F8766D", "POST" = "#00BFC4", "POST30" = "#619CFF")
pca_plot_pre_post <- pca_pre_post$plot + scale_colour_manual(values = pca_colours)
pca_plot_pre_post30 <- pca_pre_post30$plot + scale_colour_manual(values = pca_colours)

cold_pca_combined <- (pca_plot_pre_post | pca_plot_pre_post30) +
  plot_layout(widths = c(1, 1), guides = "collect") +
  plot_annotation(tag_levels = "A") &
  theme(plot.tag = element_text(face = "bold", size = 18))

ggsave(
  file.path(output_dir, "figures", "Cold_PCA_PRE_POST_and_PRE_POST30.png"),
  cold_pca_combined, width = 14, height = 6, dpi = 600, bg = "white"
)
ggsave(
  file.path(output_dir, "figures", "Cold_PCA_PRE_POST_and_PRE_POST30.pdf"),
  cold_pca_combined, width = 14, height = 6, bg = "white"
)
write.csv(pca_pre_post$scores,
          file.path(output_dir, "tables", "Cold_PCA_scores_PRE_vs_POST.csv"),
          row.names = FALSE)
write.csv(pca_pre_post30$scores,
          file.path(output_dir, "tables", "Cold_PCA_scores_PRE_vs_POST30.csv"),
          row.names = FALSE)

# ----------------------------------------------------------------------------
# 5. Cold volcano plots
# ----------------------------------------------------------------------------

prepare_volcano <- function(data) {
  data %>%
    mutate(
      Group = Regulation_nominal,
      neg_log10_p = -log10(pmax(p_value, .Machine$double.xmin))
    )
}

select_volcano_labels <- function(data, n_each = 5) {
  data %>%
    filter(Group %in% c("Up", "Down")) %>%
    group_by(Group) %>%
    arrange(p_value, desc(abs(log2FC)), .by_group = TRUE) %>%
    slice_head(n = n_each) %>%
    ungroup()
}

cold_volcano_plot <- function(data, labels, title_text, x_text,
                              common_x = NULL, common_y = NULL) {
  p <- ggplot(data, aes(log2FC, neg_log10_p, colour = Group)) +
    geom_point(size = 2.1, alpha = 0.8) +
    geom_hline(yintercept = -log10(0.05), linetype = "dashed", linewidth = 0.45) +
    geom_vline(xintercept = 0, linetype = "dashed", linewidth = 0.45) +
    geom_text_repel(
      data = labels, aes(label = Protein), size = 3.3,
      box.padding = 0.35, point.padding = 0.25,
      max.overlaps = Inf, min.segment.length = 0,
      show.legend = FALSE
```

```
    ) +
    scale_colour_manual(values = c(
      "Down" = "blue", "Not significant" = "grey70", "Up" = "red"
    )) +
    labs(
      title = title_text,
      x = x_text,
      y = expression(-log[10](p)),
      colour = NULL
    ) +
    theme_classic(base_size = 13) +
    theme(plot.title = element_text(face = "bold", hjust = 0.5))

  if (!is.null(common_x) || !is.null(common_y)) {
    p <- p + coord_cartesian(xlim = common_x, ylim = common_y)
  }
  p
}

v_post <- prepare_volcano(cold_post_vs_pre)
v_post30 <- prepare_volcano(cold_post30_vs_pre)
labels_post <- select_volcano_labels(v_post)
labels_post30 <- select_volcano_labels(v_post30)

common_volcano_x <- range(c(v_post$log2FC, v_post30$log2FC), na.rm = TRUE)
common_volcano_y <- c(0, max(v_post$neg_log10_p, v_post30$neg_log10_p, na.rm = TRUE) * 1.08)

plot_volcano_post <- cold_volcano_plot(
  v_post, labels_post,
  "Cold exposure: POST vs PRE",
  "log2 fold change (POST vs PRE)",
  common_volcano_x, common_volcano_y
)
plot_volcano_post30 <- cold_volcano_plot(
  v_post30, labels_post30,
  "Cold exposure: POST30 vs PRE",
  "log2 fold change (POST30 vs PRE)",
  common_volcano_x, common_volcano_y
)

cold_volcano_combined <- (plot_volcano_post | plot_volcano_post30) +
  plot_layout(widths = c(1, 1), guides = "collect") +
  plot_annotation(tag_levels = "A") &
  theme(plot.tag = element_text(face = "bold", size = 18))

ggsave(
  file.path(output_dir, "figures", "Cold_POST_and_POST30_volcanoes.png"),
  cold_volcano_combined, width = 14, height = 6, dpi = 600, bg = "white"
)
ggsave(
  file.path(output_dir, "figures", "Cold_POST_and_POST30_volcanoes.pdf"),
  cold_volcano_combined, width = 14, height = 6, bg = "white"
)

# ----------------------------------------------------------------------------
# 6. Exercise-cold directional comparisons
# ----------------------------------------------------------------------------

compare_exercise_with_cold <- function(exercise_data, cold_data, cold_name) {
  exercise_prepared <- exercise_data %>%
    transmute(
      Protein,
      log2FC_exercise = log2FC,
      p_exercise = p_value,
      FDR_exercise = FDR,
      ExerciseGroup = Group
    )

  cold_prepared <- cold_data %>%
    transmute(
      Protein,
      log2FC_cold = log2FC,
      p_cold = p_value,
      FDR_cold = FDR
    )

  inner_join(exercise_prepared, cold_prepared, by = "Protein") %>%
    mutate(
      ColdComparison = cold_name,
      Direction = case_when(
        log2FC_exercise > 0 & log2FC_cold > 0 ~ "Up in both",
        log2FC_exercise < 0 & log2FC_cold < 0 ~ "Down in both",
```

```
        TRUE ~ "Opposite / mixed"
      ),
      Significant_either = p_exercise < 0.05 | p_cold < 0.05,
      Significant_both = p_exercise < 0.05 & p_cold < 0.05,
      mean_abs_log2FC = (abs(log2FC_exercise) + abs(log2FC_cold)) / 2
    )
}

comparisons_post <- lapply(exercise_results, compare_exercise_with_cold,
                           cold_data = cold_post_vs_pre,
                           cold_name = "Cold POST vs PRE")
comparisons_post30 <- lapply(exercise_results, compare_exercise_with_cold,
                             cold_data = cold_post30_vs_pre,
                             cold_name = "Cold POST30 vs PRE")

comparison_post_all <- bind_rows(comparisons_post)
comparison_post30_all <- bind_rows(comparisons_post30)
comparison_all <- bind_rows(comparison_post_all, comparison_post30_all)

comparison_counts <- comparison_all %>%
  count(ColdComparison, ExerciseGroup, Direction, name = "n")

write.csv(
  comparison_all,
  file.path(output_dir, "tables", "Exercise_vs_cold_all_directional_comparisons.csv"),
  row.names = FALSE
)
write.csv(
  comparison_counts,
  file.path(output_dir, "tables", "Exercise_vs_cold_directional_overlap_counts.csv"),
  row.names = FALSE
)

# Scatter plot helper
scatter_colours <- c(
  "Down in both" = "#F8766D",
  "Opposite / mixed" = "grey70",
  "Up in both" = "#00BFC4"
)

select_scatter_labels <- function(data, n_each = 5) {
  data %>%
    filter(Direction %in% c("Up in both", "Down in both")) %>%
    group_by(Direction) %>%
    arrange(desc(Significant_either), desc(mean_abs_log2FC), .by_group = TRUE) %>%
    slice_head(n = n_each) %>%
    ungroup()
}

make_scatter <- function(data, title_text, y_text, common_x = NULL, common_y = NULL) {
  labels <- select_scatter_labels(data)
  p <- ggplot(data, aes(log2FC_exercise, log2FC_cold, colour = Direction)) +
    geom_hline(yintercept = 0, linetype = "dashed", linewidth = 0.45) +
    geom_vline(xintercept = 0, linetype = "dashed", linewidth = 0.45) +
    geom_point(size = 2.1, alpha = 0.75) +
    geom_text_repel(
      data = labels, aes(label = Protein), size = 3.1,
      max.overlaps = Inf, min.segment.length = 0,
      show.legend = FALSE
    ) +
    scale_colour_manual(values = scatter_colours) +
    labs(
      title = title_text,
      x = "Exercise log2FC POST vs PRE",
      y = y_text,
      colour = NULL
    ) +
    theme_classic(base_size = 13) +
    theme(plot.title = element_text(face = "bold", hjust = 0.5))

  if (!is.null(common_x) || !is.null(common_y)) {
    p <- p + coord_cartesian(xlim = common_x, ylim = common_y)
  }
  p
}

# Supplementary 2x2 figures for all exercise groups
make_group_scatter_set <- function(comparison_data, cold_label, y_label) {
  group_plots <- lapply(names(exercise_results), function(g) {
    d <- comparison_data %>% filter(ExerciseGroup == g)
    make_scatter(d, paste0(g, " exercise vs ", cold_label), y_label)
  })
```

```
  names(group_plots) <- names(exercise_results)
  (group_plots$Sedentary | group_plots$Endurance) /
    (group_plots$Bodybuilders | group_plots$Sprinters) +
    plot_annotation(tag_levels = "A")
}

supp_scatter_post <- make_group_scatter_set(
  comparison_post_all,
  "immediate cold exposure",
  "Cold exposure log2FC POST vs PRE"
)
supp_scatter_post30 <- make_group_scatter_set(
  comparison_post30_all,
  "cold exposure after 30 min",
  "Cold exposure log2FC POST30 vs PRE"
)

ggsave(
  file.path(output_dir, "figures", "Supplementary_exercise_groups_vs_cold_POST.png"),
  supp_scatter_post, width = 15, height = 12, dpi = 600, bg = "white"
)
ggsave(
  file.path(output_dir, "figures", "Supplementary_exercise_groups_vs_cold_POST30.png"),
  supp_scatter_post30, width = 15, height = 12, dpi = 600, bg = "white"
)

# ---------------------------------------------------------------------------
# 7. Main manuscript Figure 4
# ---------------------------------------------------------------------------

# Panel A: signed counts for the three cold contrasts
cold_count_data <- bind_rows(
  data.frame(Contrast = "POST vs PRE", Regulation = "Up", Count = 54),
  data.frame(Contrast = "POST vs PRE", Regulation = "Down", Count = -22),
  data.frame(Contrast = "POST30 vs POST", Regulation = "Up", Count = 28),
  data.frame(Contrast = "POST30 vs POST", Regulation = "Down", Count = -84),
  data.frame(Contrast = "POST30 vs PRE", Regulation = "Up", Count = 56),
  data.frame(Contrast = "POST30 vs PRE", Regulation = "Down", Count = -131)
)
cold_count_data$Contrast <- factor(
  cold_count_data$Contrast,
  levels = c("POST vs PRE", "POST30 vs POST", "POST30 vs PRE")
)

figure4A <- ggplot(cold_count_data, aes(Contrast, Count, fill = Regulation)) +
  geom_hline(yintercept = 0, linewidth = 0.5) +
  geom_col(width = 0.65, colour = "black", linewidth = 0.25) +
  geom_text(aes(label = abs(Count)),
            vjust = ifelse(cold_count_data$Count > 0, -0.3, 1.25), size = 4) +
  scale_fill_manual(values = c("Down" = "blue", "Up" = "red")) +
  scale_x_discrete(labels = c(
    "POST vs PRE" = "POST\nvs PRE",
    "POST30 vs POST" = "POST30\nvs POST",
    "POST30 vs PRE" = "POST30\nvs PRE"
  )) +
  labs(
    title = "Cold exposure-induced EV proteome remodeling",
    x = NULL,
    y = "Number of nominally altered proteins",
    fill = "Direction"
  ) +
  theme_classic(base_size = 14) +
  theme(plot.title = element_text(face = "bold", hjust = 0.5))

figure4B <- plot_volcano_post + labs(tag = NULL)

endurance_post <- comparison_post_all %>% filter(ExerciseGroup == "Endurance")
figure4C <- make_scatter(
  endurance_post,
  "Endurance exercise vs cold exposure",
  "Cold exposure log2FC POST vs PRE"
)

select_top_shared <- function(data, n_each = 6) {
  data %>%
    filter(Direction %in% c("Up in both", "Down in both")) %>%
    group_by(Direction) %>%
    arrange(desc(Significant_either), desc(mean_abs_log2FC), .by_group = TRUE) %>%
    slice_head(n = n_each) %>%
    ungroup()
}
```

```
prepare_shared_long <- function(data, cold_label = "Cold exposure") {
  up_order <- data %>% filter(Direction == "Up in both") %>%
    arrange(desc(mean_abs_log2FC)) %>% pull(Protein)
  down_order <- data %>% filter(Direction == "Down in both") %>%
    arrange(desc(mean_abs_log2FC)) %>% pull(Protein)
  protein_order <- c(rev(down_order), rev(up_order))

  data %>%
    select(Protein, Direction, log2FC_exercise, log2FC_cold) %>%
    pivot_longer(c(log2FC_exercise, log2FC_cold),
                 names_to = "Condition", values_to = "log2FC") %>%
    mutate(
      Condition = recode(
        Condition,
        log2FC_exercise = "Endurance exercise",
        log2FC_cold = cold_label
      ),
      Protein = factor(Protein, levels = protein_order)
    )
}

figure4D_selected <- select_top_shared(endurance_post, n_each = 6)
figure4D_long <- prepare_shared_long(figure4D_selected)

figure4D <- ggplot(figure4D_long, aes(log2FC, Protein, fill = Condition)) +
  geom_vline(xintercept = 0, linewidth = 0.5) +
  geom_col(position = position_dodge(width = 0.75), width = 0.65,
           colour = "grey35", linewidth = 0.25) +
  scale_fill_manual(values = c(
    "Cold exposure" = "#F8766D",
    "Endurance exercise" = "#00BFC4"
  )) +
  labs(
    title = "Directionally shared EV protein responses",
    subtitle = "Endurance exercise and cold exposure POST vs PRE",
    x = "log2 fold change", y = NULL, fill = NULL
  ) +
  theme_classic(base_size = 14) +
  theme(
    plot.title = element_text(face = "bold", hjust = 0.5),
    plot.subtitle = element_text(hjust = 0.5),
    legend.position = "bottom"
  )

figure4_main <- (figure4A | figure4B) / (figure4C | figure4D) +
  plot_annotation(tag_levels = "A") &
  theme(plot.tag = element_text(face = "bold", size = 18))

ggsave(
  file.path(output_dir, "figures", "Figure4_main.png"),
  figure4_main, width = 15, height = 11, dpi = 600, bg = "white"
)
ggsave(
  file.path(output_dir, "figures", "Figure4_main.pdf"),
  figure4_main, width = 15, height = 11, bg = "white"
)

# ---------------------------------------------------------------------------
# 8. Thesis figures: endurance comparison at both cold timepoints
# ---------------------------------------------------------------------------

endurance_post30 <- comparison_post30_all %>% filter(ExerciseGroup == "Endurance")

common_scatter_x <- range(c(
  endurance_post$log2FC_exercise,
  endurance_post30$log2FC_exercise
), na.rm = TRUE)
common_scatter_y <- range(c(
  endurance_post$log2FC_cold,
  endurance_post30$log2FC_cold
), na.rm = TRUE)

thesis_C <- make_scatter(
  endurance_post,
  "Endurance exercise vs immediate cold exposure",
  "Cold exposure log2FC POST vs PRE",
  common_scatter_x, common_scatter_y
)
thesis_D <- make_scatter(
  endurance_post30,
  "Endurance exercise vs cold exposure after 30 min",
  "Cold exposure log2FC POST30 vs PRE",
```

```
  common_scatter_x, common_scatter_y
)

thesis_cold_endurance_figure <-
  (plot_volcano_post | plot_volcano_post30) /
  (thesis_C | thesis_D) +
  plot_annotation(tag_levels = "A") &
  theme(plot.tag = element_text(face = "bold", size = 18))

ggsave(
  file.path(output_dir, "figures", "Thesis_cold_endurance_4panel.png"),
  thesis_cold_endurance_figure, width = 15, height = 11, dpi = 600, bg = "white"
)
ggsave(
  file.path(output_dir, "figures", "Thesis_cold_endurance_4panel.pdf"),
  thesis_cold_endurance_figure, width = 15, height = 11, bg = "white"
)

# Directionally shared proteins, immediate and POST30
shared_post <- select_top_shared(endurance_post, n_each = 5)
shared_post30 <- select_top_shared(endurance_post30, n_each = 5)
shared_post_long <- prepare_shared_long(shared_post, "Cold exposure")
shared_post30_long <- prepare_shared_long(shared_post30, "Cold exposure 30 min")

shared_limit <- max(abs(c(shared_post_long$log2FC, shared_post30_long$log2FC)), na.rm = TRUE)
shared_limit <- ceiling(shared_limit * 10) / 10

make_shared_plot <- function(data, subtitle_text, fill_values) {
  ggplot(data, aes(log2FC, Protein, fill = Condition)) +
    geom_vline(xintercept = 0, linewidth = 0.45) +
    geom_col(position = position_dodge(width = 0.78), width = 0.68) +
    scale_fill_manual(values = fill_values) +
    coord_cartesian(xlim = c(-shared_limit, shared_limit)) +
    labs(
      title = "Directionally shared EV protein changes",
      subtitle = subtitle_text,
      x = "log2 fold change", y = NULL, fill = "Stimulus"
    ) +
    theme_classic(base_size = 12) +
    theme(
      plot.title = element_text(face = "bold", hjust = 0.5),
      plot.subtitle = element_text(hjust = 0.5),
      legend.position = "bottom"
    )
}

shared_plot_A <- make_shared_plot(
  shared_post_long,
  "Endurance exercise and cold exposure POST vs PRE",
  c("Cold exposure" = "#F8766D", "Endurance exercise" = "#00BFC4")
)
shared_plot_B <- make_shared_plot(
  shared_post30_long,
  "Endurance exercise POST vs PRE and cold exposure POST30 vs PRE",
  c("Cold exposure 30 min" = "#F8766D", "Endurance exercise" = "#00BFC4")
)

thesis_shared_figure <- (shared_plot_A | shared_plot_B) +
  plot_annotation(tag_levels = "A") +
  plot_layout(widths = c(1, 1), guides = "collect") &
  theme(plot.tag = element_text(face = "bold", size = 18))

ggsave(
  file.path(output_dir, "figures", "Thesis_directionally_shared_proteins.png"),
  thesis_shared_figure, width = 14, height = 7, dpi = 600, bg = "white"
)
ggsave(
  file.path(output_dir, "figures", "Thesis_directionally_shared_proteins.pdf"),
  thesis_shared_figure, width = 14, height = 7, bg = "white"
)

# ---------------------------------------------------------------------------
# 9. Supplementary GO Biological Process analysis
# ---------------------------------------------------------------------------

prepare_go_input <- function(data, contrast_name) {
  data %>%
    transmute(
      Protein,
      log2FC,
      p_value,
      FDR,
```

```r
      Contrast = contrast_name,
      Regulation = classify_change(log2FC, p_value)
    )
}

go_post_input <- prepare_go_input(cold_post_vs_pre, "Cold POST vs PRE")
go_post30_input <- prepare_go_input(cold_post30_vs_pre, "Cold POST30 vs PRE")

run_go_bp <- function(result_table, direction, contrast_name) {
  query_genes <- result_table %>%
    filter(Regulation == direction) %>%
    pull(Protein) %>% unique()
  background_genes <- result_table %>% pull(Protein) %>% unique()

  if (length(query_genes) < 2) return(data.frame())

  enrichment <- gprofiler2::gost(
    query = query_genes,
    organism = "hsapiens",
    ordered_query = FALSE,
    multi_query = FALSE,
    significant = FALSE,
    exclude_iea = FALSE,
    measure_underrepresentation = FALSE,
    evcodes = FALSE,
    user_threshold = 0.05,
    correction_method = "fdr",
    domain_scope = "custom",
    custom_bg = background_genes,
    sources = "GO:BP"
  )

  if (is.null(enrichment) || is.null(enrichment$result) ||
      nrow(enrichment$result) == 0) return(data.frame())

  enrichment$result %>%
    mutate(
      Contrast = contrast_name,
      Direction = direction,
      FDR_significant = p_value < 0.05
    ) %>%
    select(any_of(c(
      "Contrast", "Direction", "source", "term_id", "term_name",
      "p_value", "FDR_significant", "term_size", "query_size",
      "intersection_size", "precision", "recall", "intersection"
    ))) %>%
    filter(FDR_significant) %>%
    arrange(p_value)
}

go_post_up <- run_go_bp(go_post_input, "Up", "Cold POST vs PRE")
go_post_down <- run_go_bp(go_post_input, "Down", "Cold POST vs PRE")
go_post30_up <- run_go_bp(go_post30_input, "Up", "Cold POST30 vs PRE")
go_post30_down <- run_go_bp(go_post30_input, "Down", "Cold POST30 vs PRE")

go_all <- bind_rows(go_post_up, go_post_down, go_post30_up, go_post30_down)
if (nrow(go_all) > 0) {
  go_all <- go_all %>%
    mutate(neg_log10_FDR = -log10(pmax(p_value, .Machine$double.xmin)))
}

write.csv(go_all,
          file.path(output_dir, "supplementary_GO", "Supplementary_GO_BP_all_results.csv"),
          row.names = FALSE)

writexl::write_xlsx(
  list(
    POST_vs_PRE_Up = as.data.frame(go_post_up),
    POST_vs_PRE_Down = as.data.frame(go_post_down),
    POST30_vs_PRE_Up = as.data.frame(go_post30_up),
    POST30_vs_PRE_Down = as.data.frame(go_post30_down),
    All_GO_results = as.data.frame(go_all)
  ),
  file.path(output_dir, "supplementary_GO", "Supplementary_GO_BP_results.xlsx")
)

# Plot significant POST30-down GO terms if present
if (nrow(go_post30_down) > 0) {
  go_plot_data <- go_post30_down %>%
    arrange(p_value) %>%
    slice_head(n = 15) %>%
    mutate(
```

```r
      neg_log10_FDR = -log10(pmax(p_value, .Machine$double.xmin)),
      term_name = factor(term_name, levels = rev(term_name))
    )

  go_plot <- ggplot(
    go_plot_data,
    aes(neg_log10_FDR, term_name, size = intersection_size,
        colour = neg_log10_FDR)
  ) +
    geom_point(alpha = 0.85) +
    labs(
      title = "Cold POST30 vs PRE: decreased proteins",
      x = expression(-log[10](FDR)),
      y = NULL,
      size = "Protein count",
      colour = expression(-log[10](FDR))
    ) +
    theme_classic(base_size = 12) +
    theme(plot.title = element_text(face = "bold", hjust = 0.5))

  ggsave(
    file.path(output_dir, "figures", "Supplementary_GO_POST30_down.png"),
    go_plot, width = 9, height = 7, dpi = 600, bg = "white"
  )
  ggsave(
    file.path(output_dir, "figures", "Supplementary_GO_POST30_down.pdf"),
    go_plot, width = 9, height = 7, bg = "white"
  )
}

# ------------------------------------------------------------------------------
# 10. Consolidated Excel workbook and session information
# ------------------------------------------------------------------------------

writexl::write_xlsx(
  list(
    Exercise_all = as.data.frame(exercise_all),
    Exercise_summary = as.data.frame(exercise_summary),
    Cold_POST_vs_PRE = as.data.frame(cold_post_vs_pre),
    Cold_POST30_vs_PRE = as.data.frame(cold_post30_vs_pre),
    Cold_POST30_vs_POST = as.data.frame(cold_post30_vs_post),
    Cold_summary = as.data.frame(cold_summary),
    Exercise_cold_comparisons = as.data.frame(comparison_all),
    Directional_counts = as.data.frame(comparison_counts),
    Figure4D_selected = as.data.frame(figure4D_selected),
    Thesis_shared_POST = as.data.frame(shared_post),
    Thesis_shared_POST30 = as.data.frame(shared_post30)
  ),
  file.path(output_dir, "EV_proteomics_complete_results.xlsx")
)

capture.output(
  sessionInfo(),
  file = file.path(output_dir, "sessionInfo.txt")
)

save.image(file.path(output_dir, "EV_proteomics_complete_workspace.RData"))

message("Analysis completed. Outputs saved to: ", normalizePath(output_dir))

# ==============================================================================
# END OF SCRIPT
# ==============================================================================
```

#### Final reporting notes

- Use “serum EV-enriched fractions” or “EV-enriched particles,” not “pure EVs.”
- The analysis is exploratory. Nominal $p < 0.05$ is used for the primary directional classification, while BH-adjusted FDR values are reported in supplementary result tables.
- Directional overlap means concordant log2FC sign and does not imply significance in both contrasts.
- The script does not include a linear mixed-effects model because the exercise analyses available here are processed group-specific result tables rather than a fully verified sample-level matrix with subject metadata.
- Before journal submission, run the script once in a clean R session and archive the resulting sessionInfo.txt, consolidated workbook, and final script together.
