## Supplementary material for "Training background shapes the exercise-induced extracellular vesicle proteome to acute exercise with partial overlap with cold exposure in humans": Suppl_Figures_and_Tables.pdf

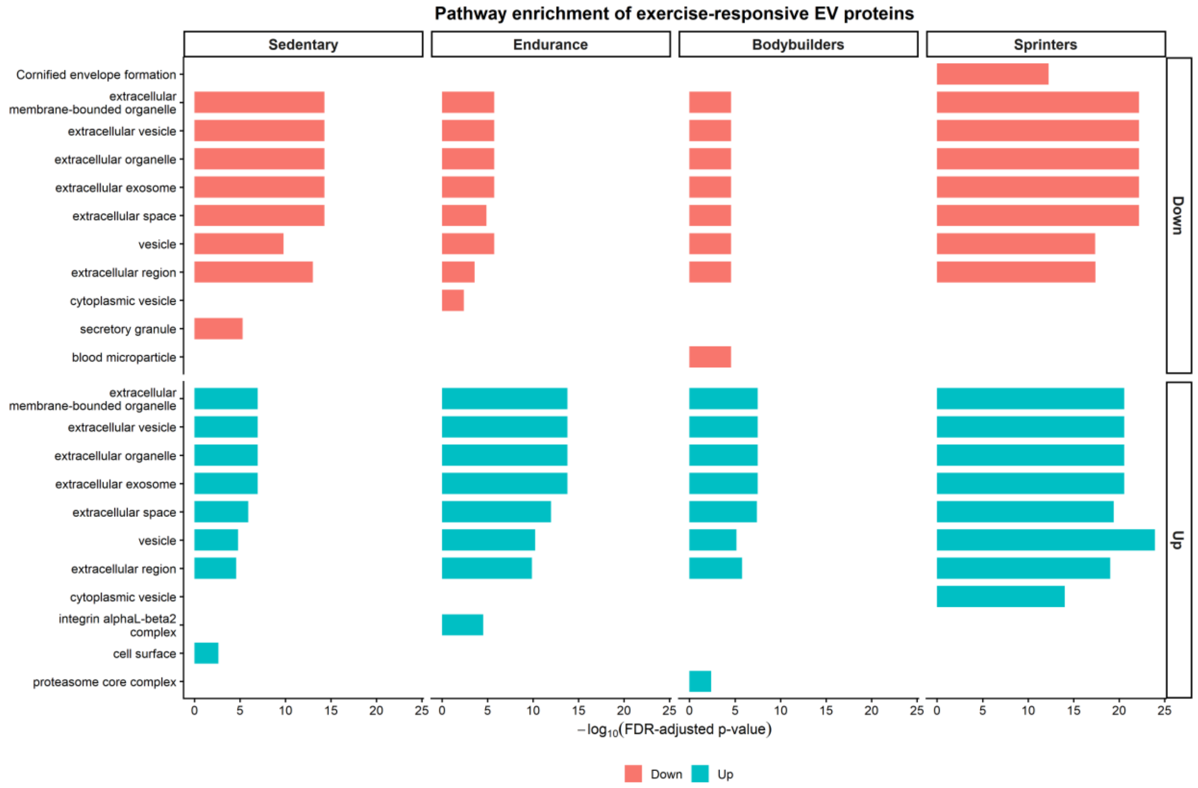


**Suppl. figure 1.** GO enrichment analysis of altered EV proteins during exercise.

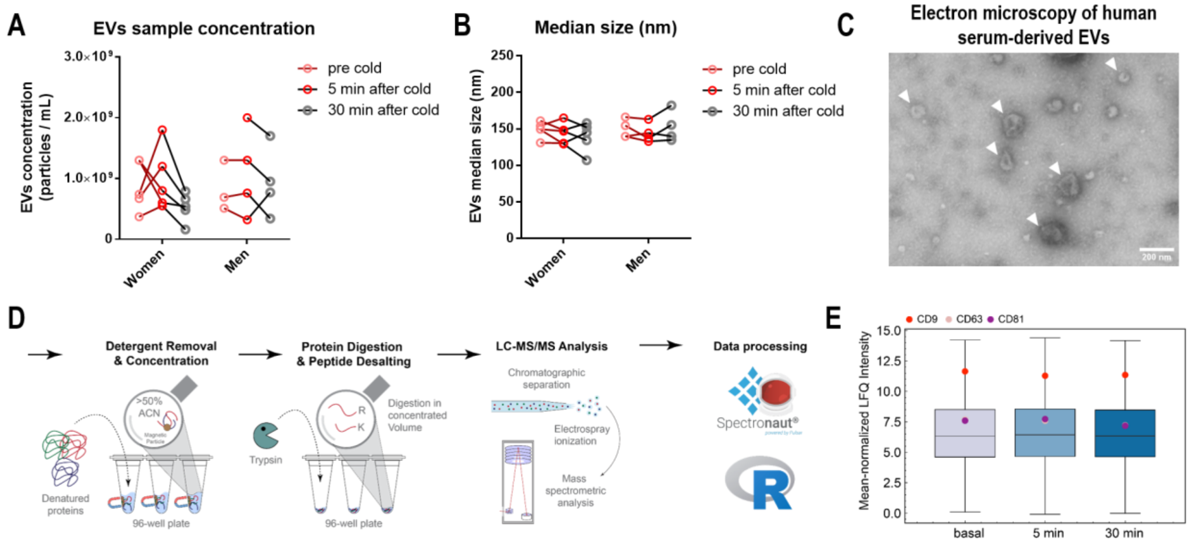


**Suppl. figure 2.** Small EVs characterization from the cold exposure intervention. Small EVs concentration (A) and median size (B) analysed by nanoparticle tracking analysis. Electron microscopy image of isolated EVs (C). Schematics showing EV proteomics workflow (D). Mean normalised Label-Free Quantification (LFQ) showing no differences in the levels of CD9, CD63 and CD81 specific markers for exosomes across the three timepoints (E).

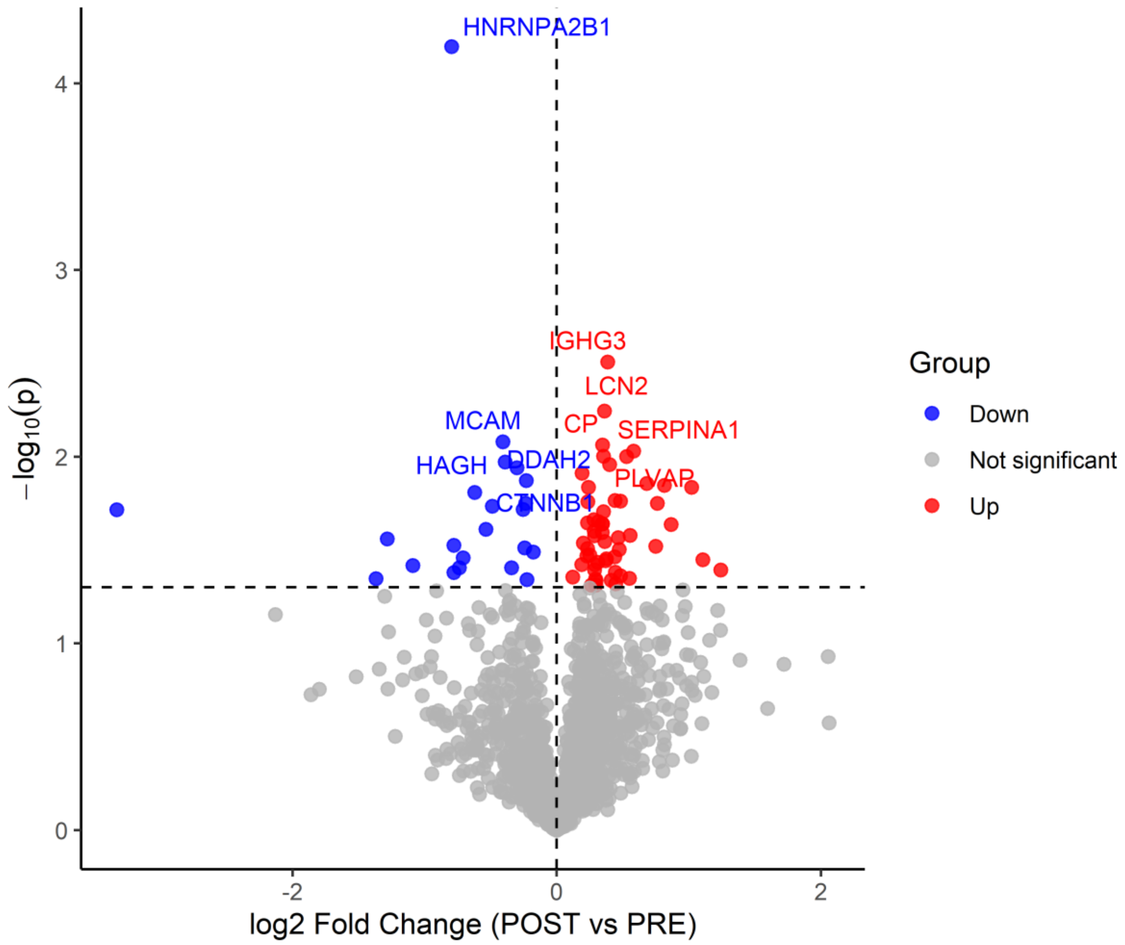


**Suppl. figure 3.** Volcano plot of significantly altered EV proteins before versus immediately after cold exposure.

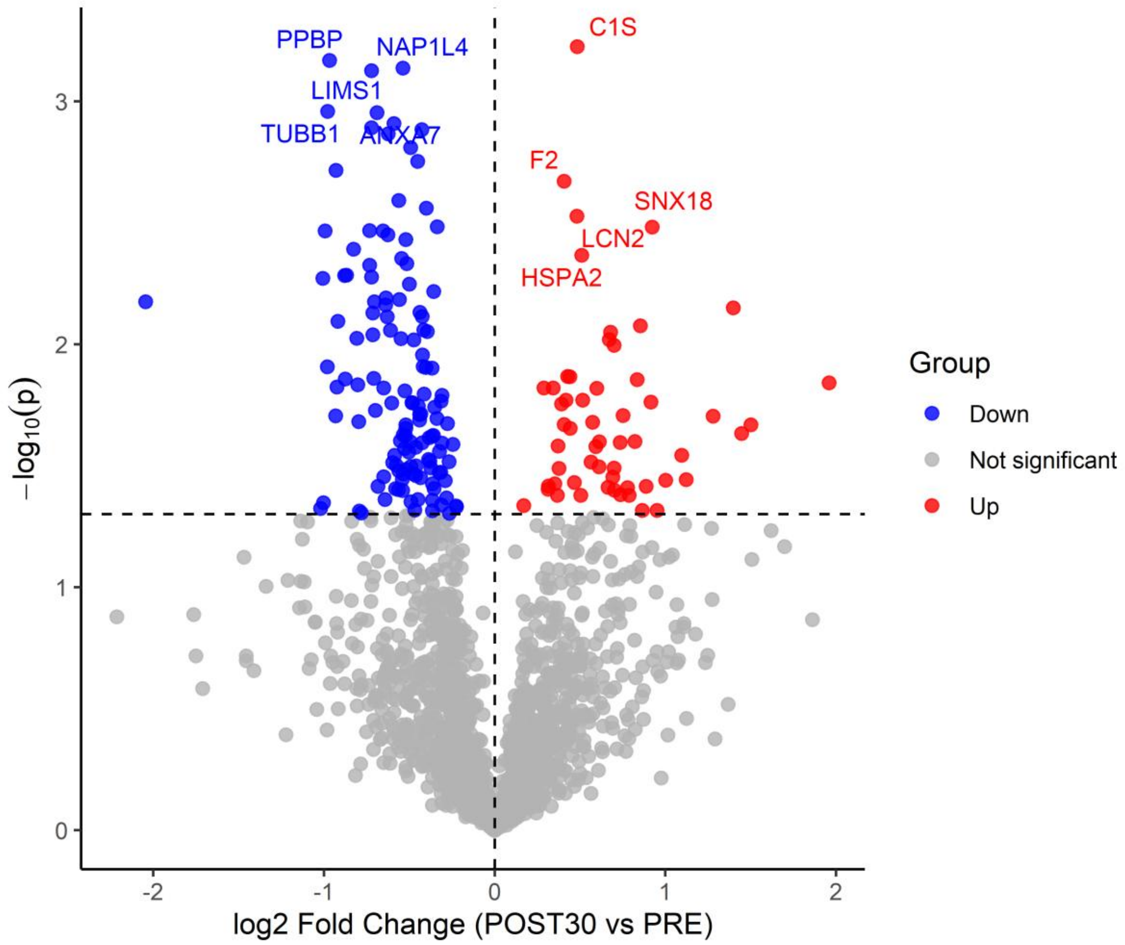


**Suppl. figure 4.** Volcano plot of significantly altered EV proteins before versus 30 minutes after cold exposure.

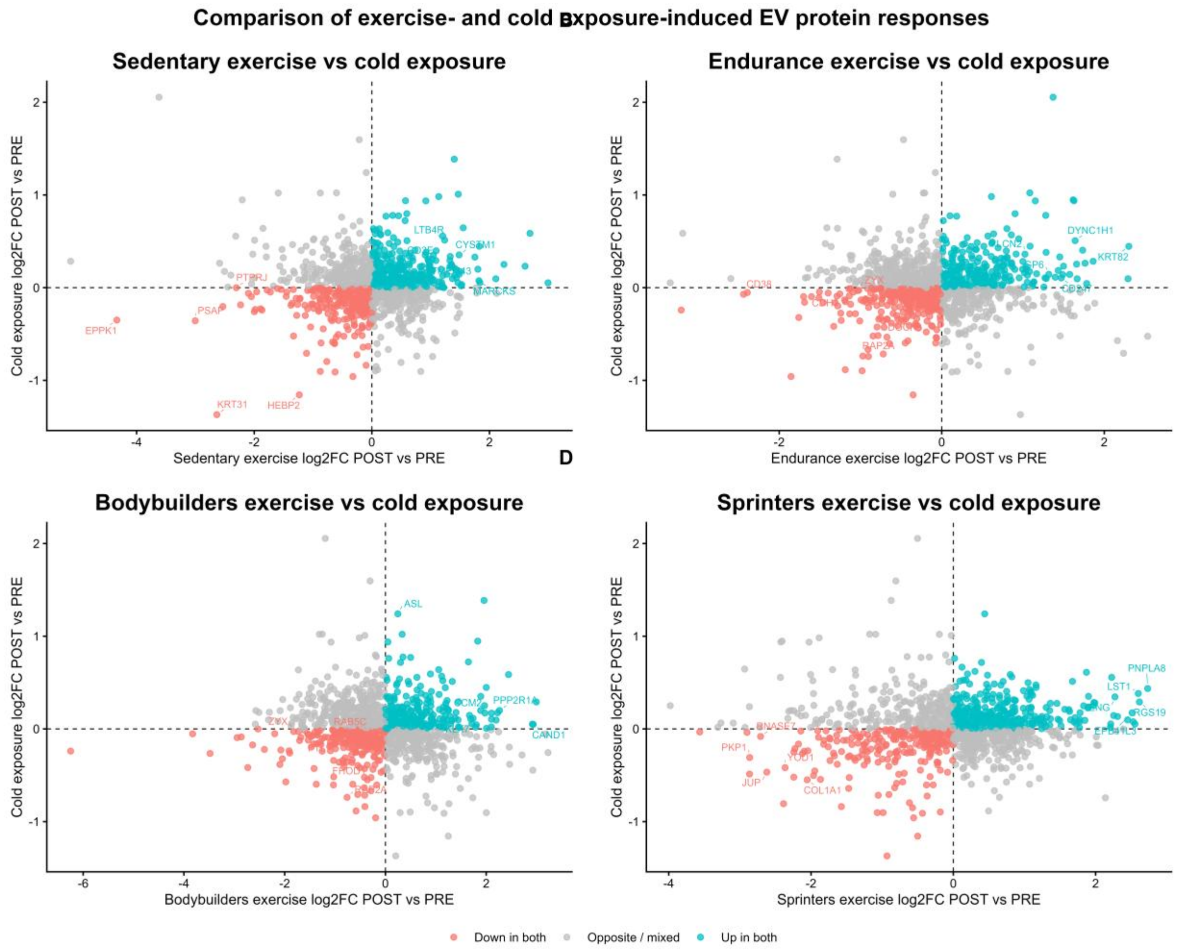


**Suppl. figure 5.** Directionally shared protein responses between cold POST vs PRE cold exposure and POST vs PRE exercise in sedentary controls, endurance athletes, bodybuilders and sprinters.

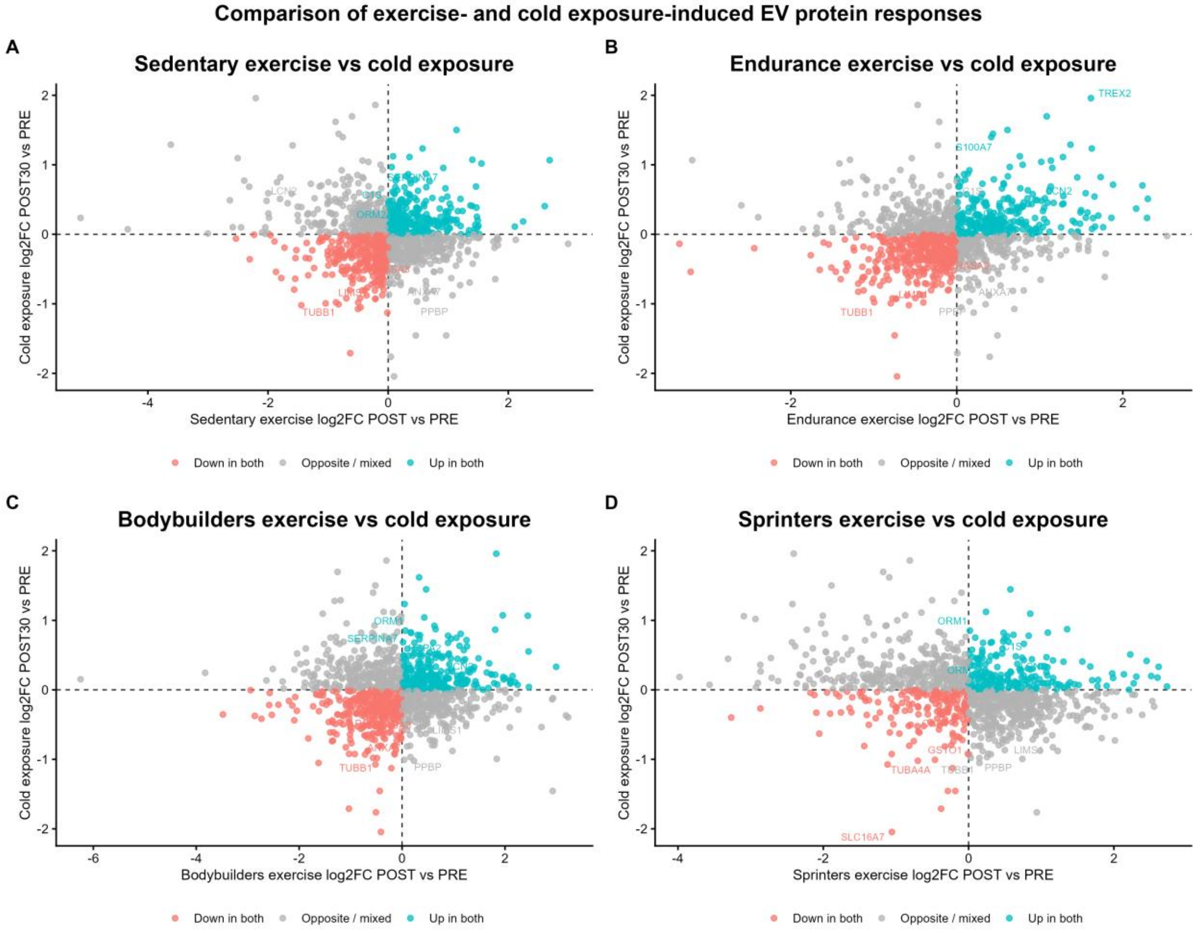


**Suppl. figure 6.** Directionally shared protein responses between cold POST30 vs PRE cold exposure and POST vs PRE exercise in sedentary controls, endurance athletes, bodybuilders and sprinters.
